# Visual integration of omics data to improve 3D models of fungal chromosomes

**DOI:** 10.1101/2023.03.28.534549

**Authors:** Thibault Poinsignon, Mélina Gallopin, Pierre Grognet, Fabienne Malagnac, Gaëlle Lelandais, Pierre Poulain

## Abstract

The functions of eukaryotic chromosomes and their spatial architecture in the nucleus are reciprocally dependent. Hi-C experiments are routinely used to study chromosome 3D organization by probing chromatin interactions. Standard representation of the data has relied on contact maps that show the frequency of interactions between parts of the genome. In parallel, it has become easier to build 3D models of the entire genome based on the same Hi-C data, and thus benefit from the methodology and visualization tools developed for structural biology. 3D modeling of entire genomes leverages the understanding of their spatial organization. However, this opportunity for original and insightful modeling is under exploited. In this paper, we show how seeing the spatial organization of chromosomes can bring new perspectives to Hi-C data analysis. We assembled state-of-the-art tools into a workflow that goes from Hi-C raw data to fully annotated 3D models and we re-analysed public Hi-C datasets available for three fungal species. Besides the well-described properties of the spatial organization of their chromosomes (Rabl conformation, hypercoiling and chromosome territories), our 3D models highlighted *i)* in *Saccharomyces cerevisiae*, the backbones of the cohesin anchor regions, which were aligned all along the chromosomes, *ii)* in *Schizosaccharomyces pombe*, the oscillations of the coiling of chromosome arms throughout the cell cycle and *iii)* in *Neurospora crassa*, the massive relocalization of histone marks in mutants of heterochromatin regulators. 3D modeling of the chromosomes brings new opportunities for visual integration. This holistic perspective supports intuition and lays the foundation for building new concepts.

## Introduction

What if it were possible to see all the details of chromosomes inside the nucleus of a cell? In eukaryotic cells, the nucleus is a dynamic organelle which is highly organized and characterized by extensive compartmentalization of structural components in its three-dimensional space (see (Razin et al. 2014; Dundr and Misteli 2001; Cremer and Cremer 2001; Arifulin et al. 2018) for reviews). In such a crowded environment, the arrangement of chromosomes is constrained and requires the formation of multiple chromatin domains to limit gene positions to preferred locations within the nuclear space (Misteli 2020). The spatial organization of chromosomes is of great interest, helping molecular biologists represent the objects they work with, and understand their interactions. Even if immense progress has been made in cell imaging, biological molecules (e.g. DNA, RNA, proteins) are too small to be individualized with optical microscopes (Wong and Eleftheriades 2013; Jensen 2013) and consequently, the interior of the cell (and the interior of a nucleus even more) remains largely invisible to the human eye. Alternative solutions are based on molecular-scale techniques like X-ray crystallography (Smyth 2000), NMR microscopy (Reckel et al. 2005) or electron microscopy (Radulović et al. 2022). By analyzing atom arrangements in molecules, these techniques produce informative views of macromolecular complexity (Nogales and Scheres 2015; Baumeister 2022), but they require complex technical skills and expensive equipment. Furthermore, it is important to keep in mind that in the end, these images are still artificial representations of reality. In other words, they are “models”.

The use of models is widespread in biology. From model organisms to model systems, their interest is to understand a phenomenon in a simplified context, in order to, later, generalize to more complex situations. In cell biology for instance, structural models are used to represent cell components (membranes, nucleus, cytoplasm, etc.), to understand their organization, and to describe their constituent molecules (Im et al. 2016). The work of David Goodsell provides an emblematic example (Goodsell 2009). His drawings representing cellular compartments and their molecular actors are so striking because of the unexpected density of molecules and the complexity of their organization. Goodsell’s illustrations have been featured as “Molecule of the Month’’ on the Protein Data Bank website for over twenty years and the scientific journal Nature chose one of his paintings to make the cover of a special issue on COVID-19 (August 20, 2020 issue). Models make it possible to represent and summarize, in an intuitive but still scientifically rigorous way, the massive knowledge of cell molecular structures. Creating them thus represents a stimulating challenge, at the crossroads of multiple disciplines (biology, physics, computer science, art) (O’Donoghue 2021).

Modeling chromosomes is challenging because they belong to the mesoscale, *i. e.* a length-scale that is larger than discrete molecular complexes yet still remains intracellular (Sear et al. 2015). As a consequence, they are both too small to be observed with precision under optical microscopes, and too large to be fully modeled at the atomic scale. In eukaryotes, chromosomes are long DNA molecules, tightly packed to fit within the nucleus, a space only a fraction of their length. To this end, DNA is wrapped around histone proteins to form nucleosomes, stacked to form chromatin fibers, themselves arranged into higher-order chromatin architecture (Misteli 2020). In this study, we are specifically interested in seeing the spatial organization of fungal chromosomes. Our laboratory has long-standing expertise in functional genomics projects in yeasts (Denecker et al. 2020; Poinsignon et al. 2022; Sénécaut et al. 2022) or filamentous fungi (Grognet et al. 2019; Carlier et al. 2021; Lelandais et al. 2022), and spatial genome organization in fungi has already been investigated, for model species like *Saccharomyces cerevisiae* (Duan et al. 2010; Tokuda et al. 2012), *Schizosaccharomyces pombe* (Grand et al. 2014; Tanizawa et al. 2010; Gallardo et al. 2019; Noma 2017) and *Neurospora crassa* (Galazka et al. 2016; Rodriguez et al. 2022). Notably, the nuclear architecture of *N. crassa*, a multicellular fungus that grows as a mycelium with a network of hyphae (Galagan et al. 2003), has structural homology (thanks to the existence of heterochromatin and euchromatin) with the human genome (Rodriguez et al. 2022). This makes the *N. crassa* genome a cost-efficient model to study chromosome conformation. *S. cerevisiae* and *S. pombe* are distantly related yeast species (evolutionary distance of at least 400 Mya) that represent very different models of unicellular eukaryotes. Their genomes, of similar size (∼12 Mb), are organized into different sets of chromosomes (16 chromosomes for *S. cerevisiae* and 3 chromosomes for *S. pombe*). These yeast genomes are more than three times shorter than the genome of *N. crassa* (genome size is 41 Mb, with 7 chromosomes ranging from 4 to 10 Mb). Altogether, *N. crassa*, *S. cerevisiae* and *S. pombe* represent an interesting diversity of genomic situations for which much knowledge of nuclear organization is available in the literature (to view representative data from several publications, see **Supplementary Figure S1**).

The most emblematic feature highlighted in these articles is the “Rabl conformation”. This is a particular positioning of chromosomes in which their centromeres and telomeres are respectively clustered at distinct peripheral locations, inside the nuclear envelope. The Rabl conformation has been observed in *N. crassa*, *S. cerevisiae* and *S. pombe* (**Supplementary Figure S1**). It has been found in plants (Rodriguez-Granados et al. 2016), mice (Stevens et al. 2017; Zhang et al. 2020) and flies (Bauer et al. 2012), underpinning the idea that it represents a conserved constrained genome structure to limit topological entanglement of chromosomes (Pouokam et al. 2019). The clustering of centromeres and telomeres at distinct peripheral locations inside the nuclear envelope induces chromosomes to be organized in a “clothespin-like” structure, meaning that they are folded in half at the centromere, their arms against each other, their ends slightly bent. In *S. pombe*, the clustering of centromeres and telomeres is reinforced by protein anchors to the nuclear envelope and heterochromatin epigenetic marks (**Supplementary Figure S1**). At the molecular level, 3D genome organization in yeasts is determined by cohesin and condensin mediated structures (36,37). In yeasts, cohesin seems to form 40∼50 kb globules (Mizuguchi et al. 2014; Tanizawa et al. 2017), and condensin 300∼500 kb large domains (Noma 2017; Tanizawa et al. 2017; Kim et al. 2016). This highlights how 3D genome structures are organized at successive, nested scales of condensation patterns. Another important feature relates to the definition of “chromosome territories’’, where chromosomes are folded within their own space in the nucleus (Fritz et al. 2019). Those territories are another well described example of the functional importance of chromatin structure and are observed in many different organisms. Well known in yeasts, chromosome territories are still debated in *N. crassa* (Rodriguez et al. 2022). Nonetheless correlations between 3D structuration and heterochromatin/euchromatin epigenetic marks were revealed by Hi-C and ChIP-seq studies (Rodriguez et al. 2022; Pouokam et al. 2019; Fritz et al. 2019). Centromeres and telomeres are heterochromatic (as in yeasts) and euchromatic regions are condensed into 20∼40 kb globules (consistent with yeast cohesin globules).

This knowledge of the spatial organization of *N. crassa*, *S. cerevisiae* and *S. pombe* chromosomes was mainly built on Hi-C data analyses. The initial 3C technique evolved rapidly with NGS technologies, and recent technical improvements allow single cell analysis, with better spatial resolution (Hsieh et al. 2016) or haploid investigations (Oomen et al. 2020; Mitter et al. 2020) (see (Kempfer and Pombo 2020; Jerković and Cavalli 2021) for reviews). Various types of Hi-C data are thus available in public databases, differing in accuracy and resolution, for numerous species (Hoencamp et al. 2022), far beyond fungal models. Their processing steps are now well described (see (Jerković and Cavalli 2021) for review), and bioinformatics tools like the Juicer suite (Durand et al. 2016) or the HiC-Pro pipeline (Servant et al. 2015) are available and widely used by the community. Generally, the analysis of Hi-C data ends with the representation of a “contact map” (symmetrical heatmap). A contact map shows the frequency of contacts observed between different portions of a genome, whose sizes depend on a predefined resolution (generally around 5 to 10 kb). Contact maps present the advantage of summarizing in a single image, all pairwise distances between genomic regions, both at short and long-range. Contact maps thus reveal chromosome clusters, their organization into domains and the formation of DNA loops. They are valuable because of their precision (resolution can go down to a few kb) and the information density they convey. Still, their biological interpretation is not trivial and, in the end, contact maps remain abstract representations of the spatial organization of chromosomes.

3D modeling of Hi-C contact maps is an interesting strategy to boost interpretation of Hi-C experimental results (Yardımcı and Noble 2017; Oluwadare et al. 2019). It consists in calculating 3D coordinates (x, y, z) for all genomic intervals shown in a contact map, so that their pairwise Euclidean distances remain consistent with their contact frequencies in the original map. 3D modeling of the spatial organization of chromosomes is not trivial, but several software packages based on different strategies exist. For instance, “distance-based methods” convert the number of contacts stored in the contact maps into distances and then resolve an optimization problem to fit 3D coordinates to those distances (Rieber and Mahony 2017; Li et al. 2018). Such methods often require fulfilling physical constraints related to known structural features of the genomes (for instance centromere and telomere clustering in opposite locations inside the nucleus or rDNA clustering outside the overall structure to form the nucleolus). This is how the 3D models already proposed for *N. crassa*, *S. cerevisiae* and *S. pombe* were obtained (**Supplementary Figure S1**, black stars). Alternatively, “probability-based strategies” model contact numbers using random variables. This offers the advantage of taking into account the nature of Hi-C measurements (an average over a cell population) (Varoquaux et al. 2014). As an illustration, the Pastis-NB method (implemented in the Pastis software (Varoquaux et al. 2021)) implements a probability-based strategy based on negative binomial random variables. This distribution properly models count data generated by sequencing technologies, as illustrated by the wide adoption of negative binomial distributions to model RNA-seq data (Love et al. 2014). The model therefore accounts for over-dispersion, performs well in low coverage settings and doesn’t require any initial physical constraints. Once a 3D model has been generated, a large panel of tools exists to visualize the 3D coordinates of genome models: Genome3D, GMOL, GenomeFlow, HiC-3DViewer, Csynth, WashU Epigenome Browser (Asbury et al. 2010; Nowotny et al. 2016; Trieu et al. 2019; Djekidel et al. 2017; Todd et al. 2021; Li et al. 2022). Initially limited to the context of viewing biomolecules (nanoscale), visualization software was thus extended to representation of the 3D organization of chromosomes at the mesoscale.

Surprisingly, recent Hi-C studies on *S. cerevisiae*, *S. pombe* and *N. crassa* do not feature 3D models (Rodriguez et al. 2022; Tanizawa et al. 2017; Costantino et al. 2020). In a context where *i*) an increasing number of Hi-C studies are generated, *ii)* methods for 3D modeling of Hi-C contacts exist and *iii)* software for 3D visualization is routinely available in structural biology, one may wonder why contact maps are not associated with 3D models more often. In the present study, we explored the potential of using 3D models as well as Hi-C contact maps to better understand the spatial organization of fungal chromosomes. For this purpose, we created a workflow called 3DGB to simplify the creation of 3D models from Hi-C data. Open source and freely available on GitHub, 3DGB generates contact maps, builds 3D models and adds further processing of the 3D model output as PDB files, suitable for advanced visualization with molecular viewer software. Using 3DGB, we created several models of the *S. cerevisiae*, *S. pombe* and *N. crassa* genomes, starting from Hi-C data available in public databases. Different strains were analyzed (wild type and mutants for heterochromatin organization or structural proteins), at different stages of the life cycle. Our models showcase known characteristics of the chromosomal organization of these genomes. But they also reveal, thanks to the visual integration of omics data, important properties of regulatory proteins with critical functions for the maintenance of spatial organization. We thus demonstrate the interest of 3D modeling of Hi-C contacts for studies of genome organization.

## Results

### Part 1. Simplifying 3D model creation for Hi-C data analysts with a robust and reproducible workflow

#### General overview of 3D Genome Builder

To simplify the creation of 3D models for researchers interested in Hi-C data analysis, we created 3D Genome Builder (3DGB), a bioinformatics workflow that streamlines the generation of 3D models to visualize and explore the spatial organization of chromosomes, based on Hi-C experimental results. A general overview of the workflow is presented in **Figure 1**. Starting from Hi-C raw FASTQ files, 3DGB automatically performs the critical bioinformatics steps required to *i)* compute Hi-C contact frequencies, *ii)* infer associated 3D models of the chromatin organization and *iii*) annotate and control the quality of the 3D models. These 3D models are stored in standard PDB files, so they can be further investigated with complementary visualization tools (see below). 3D models can optionally be enriched with supplementary quantitative omics data, such as ChIP-seq or RNA-seq signals (see next section). 3DGB has been designed to remain as simple as possible. Therefore, it requires only four basic inputs to be specified by the user (**Figure 1**, green boxes): FASTQ file identifiers, a FASTA file with the sequence of the reference genome, the list of restriction sites for the enzymes used during the Hi-C experiments, and the targeted resolutions for the Hi-C data analysis. This final parameter has an impact on the final 3D models, *i.e.* the smaller the value (specified in bp), the more detailed the 3D model.

**Figure 1:**
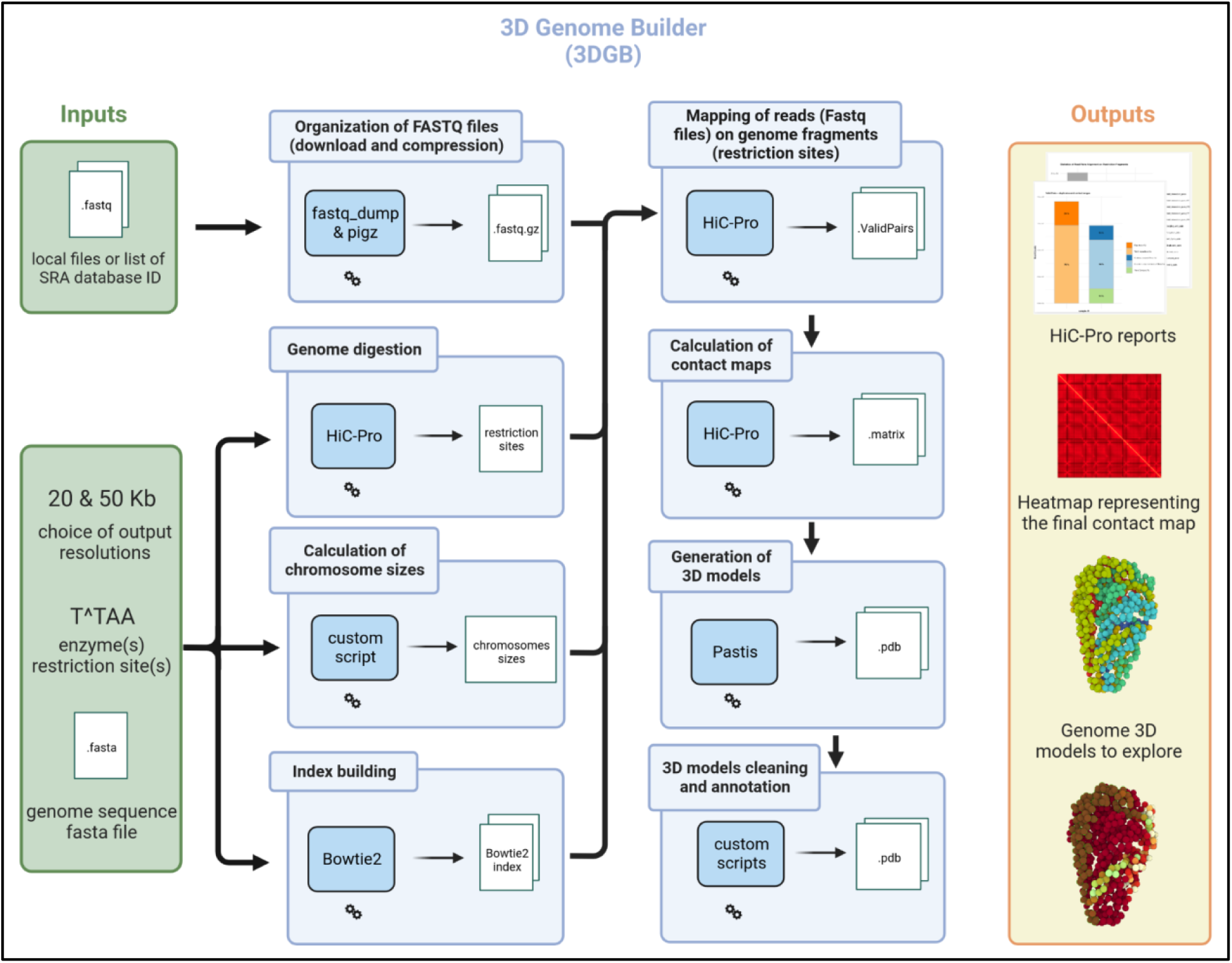
General overview of the 3D Genome Builder (3DGB) workflow. Inputs and outputs are respectively represented with green and orange boxes (left and right panels). The workflow is detailed as a sequence of blue boxes (middle panel). Each blue box represents a task of particular interest. The 3DGB workflow is orchestrated with Snakemake (see **Supplementary Data S2** and **Methods**). All tasks are automatically chained one after the other until target outputs are produced. Final results comprise quality control reports generated by HiC-Pro, contact heatmaps and PDB files with the 3D models. PDB files are enriched with additional annotations (see main text for more details) and can be visualized with any PDB viewer software (mol* in our case).

The eight main steps required for Hi-C data processing are represented in blue boxes in **Figure 1** and rely on two state-of-the-art software packages. The first step utilized HiC-Pro (Servant et al. 2015), a reference in Hi-C data processing, cited more than 800 times. It processes raw FASTQ files, performs quality control and generates normalized contact counts and associated figures (presenting important statistics to evaluate the quality of read mapping and justify potential read filtering). Then, Pastis (Varoquaux et al. 2021) iteratively computes 3D models of the organization of chromosomes, through an original negative binomial contact count modelization (referred to as Pastis-NB). Interestingly, it outputs a consensus model, which, by its uniqueness, greatly simplifies downstream analyses and interpretations. Around those two main components (HiC-Pro and Pastis), 3DGB centralizes the configuration, performs contact calculations, generates contact heatmaps, builds 3D models and adds further processing of the 3D model output as PDB files. 3DGB is open source, available in GitHub (https://github.com/data-fun/3d-genome-builder) and archived in Software Heritage ([swh:1:dir:26b6504724952e6d0d7db34c394e052217523754](https://archive.softwareheritage.org/browse/directory/26b6504724952e6d0d7db34c394e052217523754/?origin_url=https://github.com/data-fun/3d-genome-builder&revision=898faf98dfb66d53e7f3e9934c9258fcf0abfacd&snapshot=7d4cd336f896654b665fc9f837d00fbaca54ad2a)).

#### Enriched output PDB files for advanced visualization with molecular viewer software

The main 3DGB outputs are 3D models of genome organization. These 3D models are composed of beads for which 3D coordinates (x, y, z) were inferred from contact information (see **Methods**). One bead represents several thousand base pairs corresponding to the chosen resolution during the Hi-C data analysis (see previous section). To facilitate the study of these models, we enriched the structures produced by Pastis as PDB files with a four-step procedure. First, 3DGB formats and enriches the 3D model output by Pastis for visualization and data integration by annotating each bead with the chromosome number. This is useful to distinguish chromosomes when viewing complete structures (see **Figure 2** for illustrations). Second, it automatically reconstructs beads for which no coordinates could be calculated by Pastis, by interpolating missing coordinates from the existing ones (see **Methods**). Note that we do not extrapolate missing coordinates, meaning that beads with missing coordinates located at the extremities of the model are discarded. Third, outlier beads, *i.e.* beads placed outside the overall model, are filtered out and deleted based on a threshold value which can be specified by the user. Four, quantitative values, such as ChIP-seq data, can be used to color the model, allowing visual integration of omics data on the 3D structure of the genome (see next section for detailed examples of such integration of omics data). All these additional functionalities were implemented in Python (Guido Van Rossum and Fred L. Drake 2009) scripts integrated within the Snakemake workflow (see **Methods)**.

**Figure 2:**
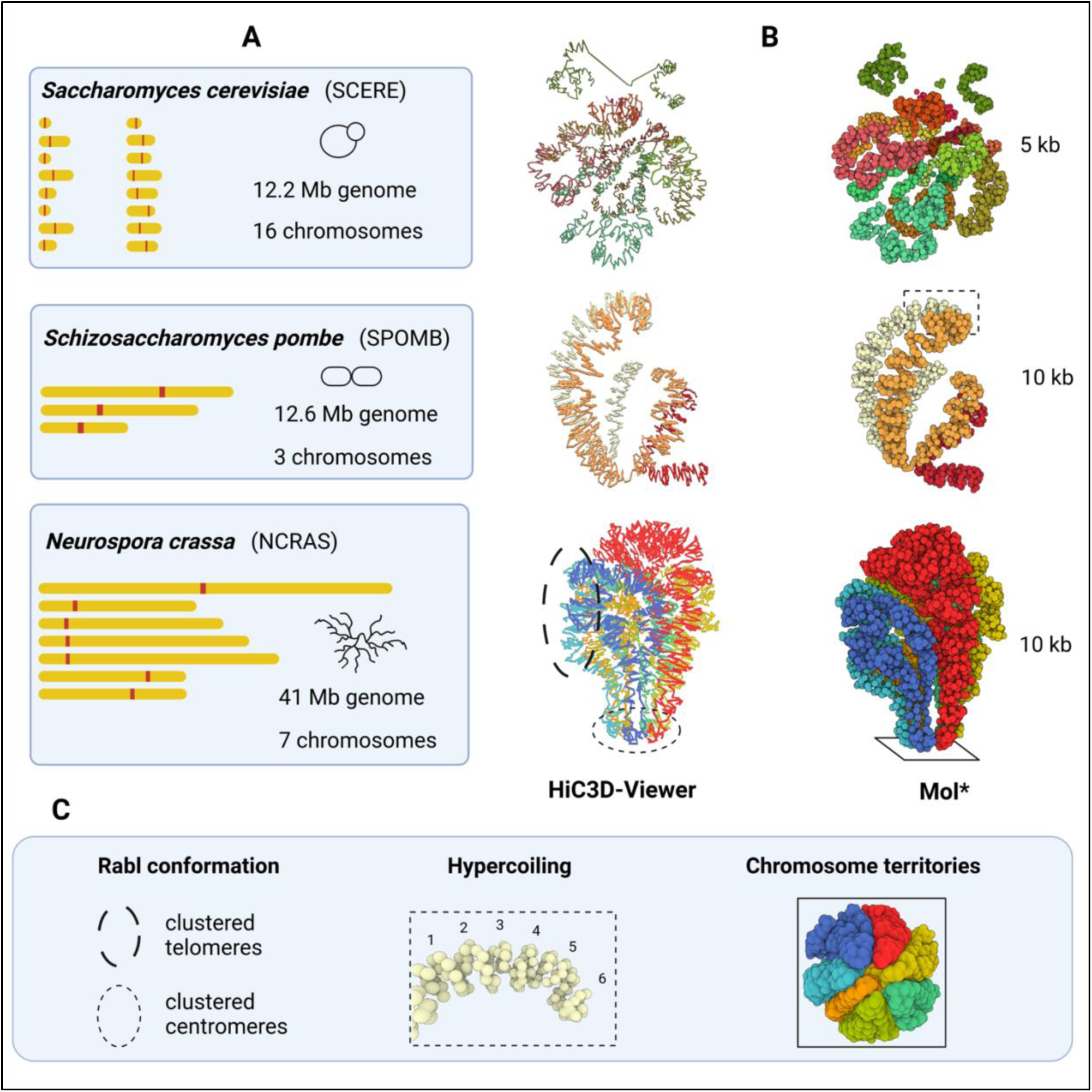
3D modeling of fungal chromosomes in wild type situations, in the model yeasts S. cerevisiae and S. pombe and the filamentous fungus N. crassa. **(A)** 2D representations of chromosomes in each species. They are displayed with the same scale, showing the diversity of genomes used in this study to create 3D models. (B) 3D models obtained with the 3DGB workflow. The source of the Hi-C data used to create them are presented in Table 1. They are visualized with HiC3D-Viewer (left) and Mol* (right). Note that HiC3D-Viewer links beads artificially, whereas with Mol*, beads remain individualized (they each represent a 5 or 10 kb chromatin region, depending on the species, see Table 1). (C) Key features of genome organization are highlighted. The Rabl configuration, the hypercoiling of chromosomes and their distribution into separate territories are well observed on the 3D models obtained with 3DGB and presented in (B) (see circles and rectangles)

Altogether, these enhancements provide convenient 3D models to be viewed and explored by Hi-C data analysts. For chromatin 3D model file formats, we provided the 3D model structure in the PDB format, a traditional file format widely used to store coordinates of molecular structures. Most software used in structural biology to view and manipulate structures can handle the PDB format. In this paper, we preferentially used Mol* (Sehnal et al. 2021), a ubiquitous viewer for large scale molecular structures, with a user-friendly web interface allowing visualization and customization of 3D models with only a few mouse clicks. We also used HiC3D-Viewer (Djekidel et al. 2017), a viewer especially developed for chromatin structures. Note that HiC3D-viewer requires conversion of the PDB file format into the G3D format. 3DGB also provides 3D models in the G3D file format, allowing users to choose the viewing software they prefer. As illustrations, images produced by Mol* and HiC3D-viewer are presented in **Figure 2**, for three different 3DGB models inferred in yeasts and a filamentous fungus (see next section).

### Part 2. Seeing the spatial organization of fungal chromosomes with 3D Genome Builder (3DGB)

#### Creation of wild type models and visual consistency with the literature

To test the performances and the relevance of 3DGB, we chose three emblematic fungal species: *Saccharomyces cerevisiae*, *Schizosaccharomyces pombe* and *Neurospora crassa*. We wanted to create 3D models of their genome organization and confront the models with current knowledge found in the literature. The *S. cerevisiae* Hi-C data are from a recent study from Constantino et al. (Costantino et al. 2020), the *S. pombe* Hi-C data are from Tanizawa et al. (Tanizawa et al. 2017) and the *N. crassa* Hi-C data are from Galaska et al. (Galazka et al. 2016). Detailed information regarding the source of the data are given in **Table 1**. As a result, we obtained new models of *S. cerevisiae*, *S. pombe* and *N. crassa* genomes. Three of them, which correspond to wild type situations, are shown in **Figure 2**. As expected, the key features of the spatial organization of the three species’ genomes were observed, *i.e.* the Rabl conformation, the hypercoiling of chromatin fibers and the chromosome territories. Finding these well-described characteristics was an important step in validating the relevance of the 3DGB workflow, especially considering that 3DGB uses a probabilistic-based strategy to infer 3D models (Pastis-NB), which is free of initial constraints, such as the relative position of chromosomes.

**Table 1:**
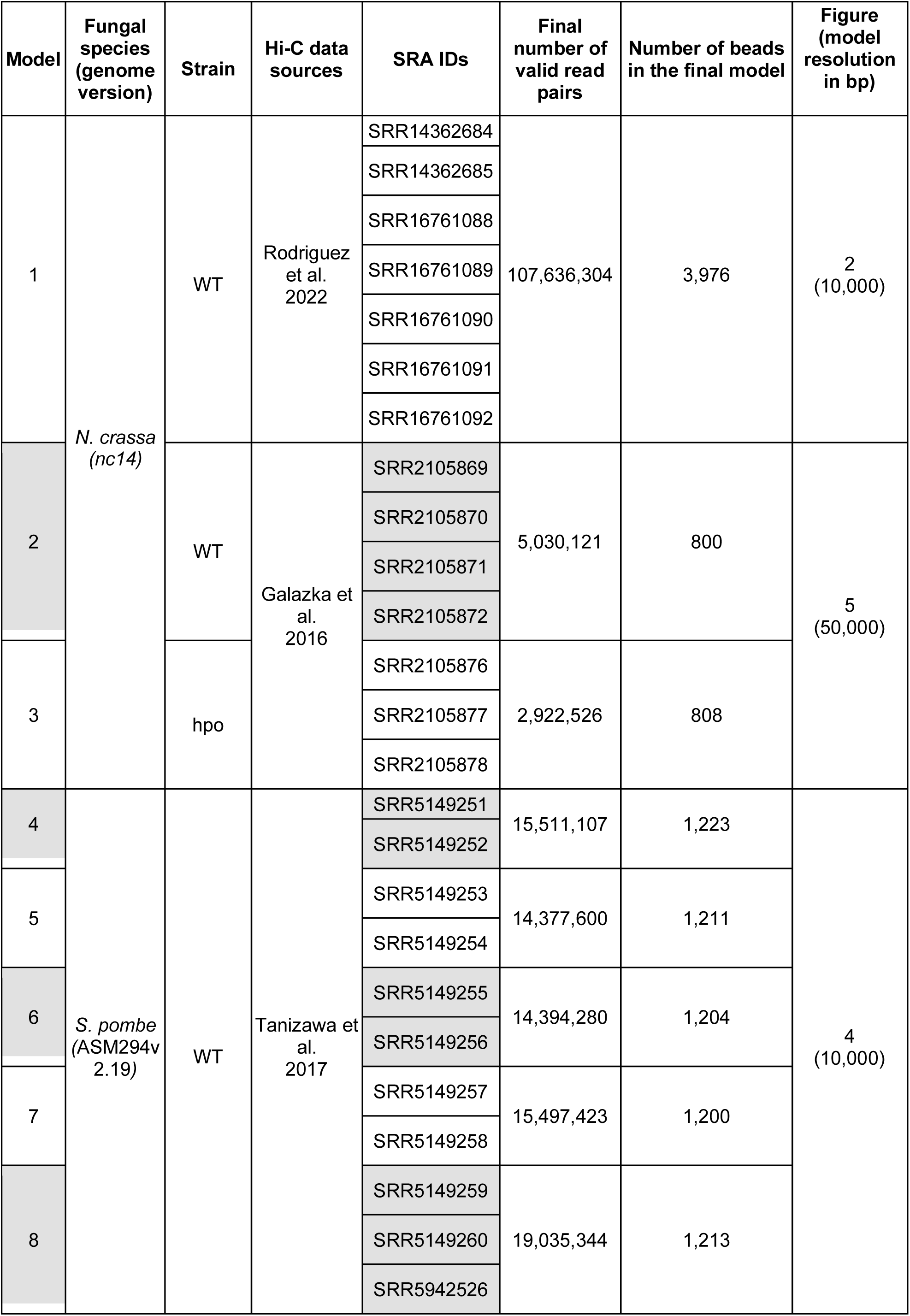
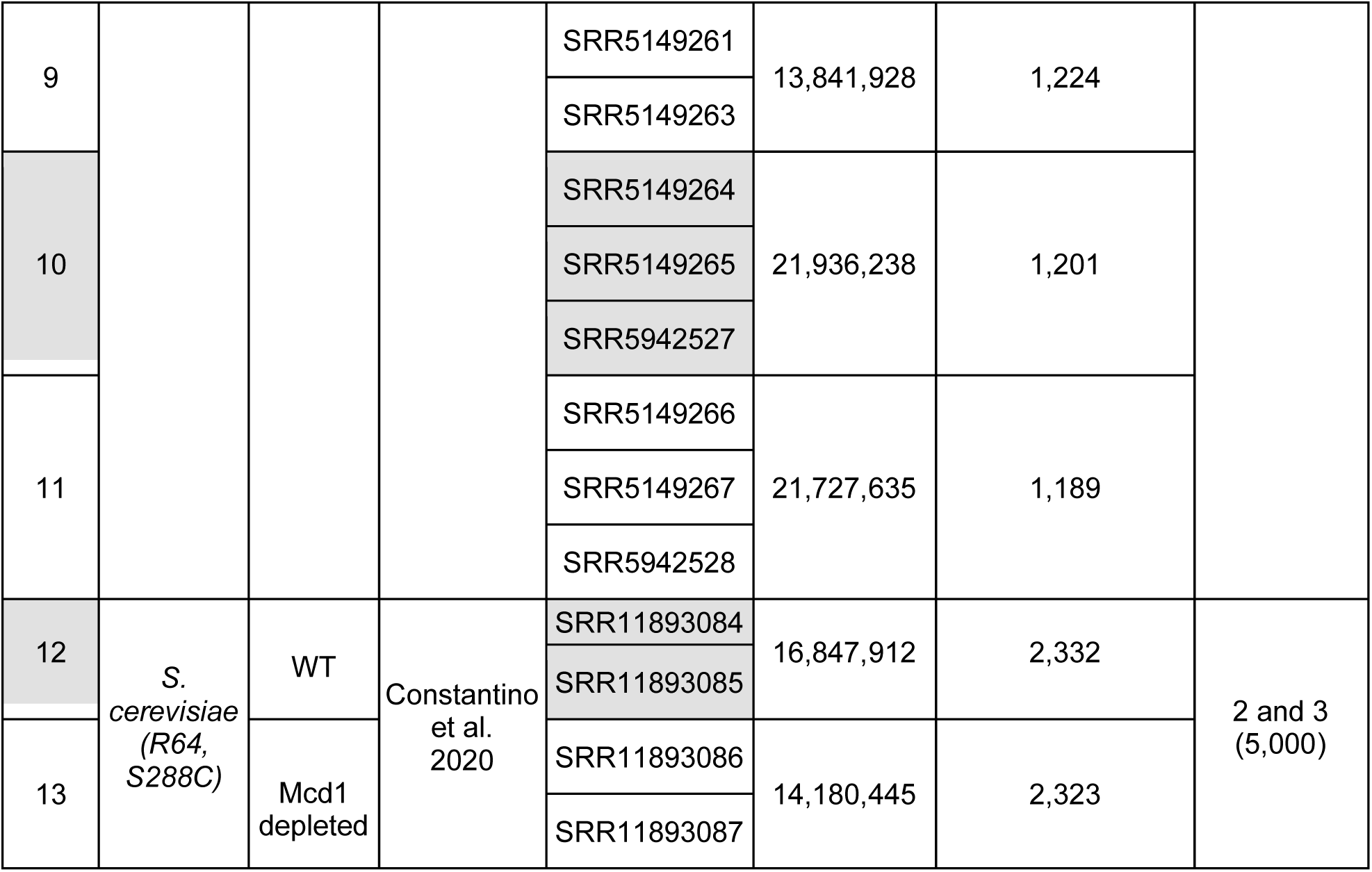
Main characteristics of raw data used in this study to create 3D models of fungal chromosomes. The sources of Hi-C data (original articles and SRA identifiers) are given. All datasets were obtained on wild type strains, with the exception of models #3 and #13 which correspond to the N. crassa hpo mutant (used in **Figure 5**) and S. cerevisiae mcd1 depleted (used in **Figure 3**). The number of valid pairs of reads gives an estimation of the overall quality of the data and indicates the density of the contact frequencies measured in the experiment. The number of beads in the 3D model is determined by the chosen resolution during Hi-C data analysis and the length of the reference genome.

#### Added value of 3DGB models for visualization: examples ranging from a detailed view to a more general view of chromatin organization

##### Example #1: Spatial alignment of cohesin binding sites along chromosomes in *S. cerevisiae*

Ten years after the first 3D model of the *S. cerevisiae* genome (see (Duan et al. 2010) and **Supplementary Figure S1**), Costantino et al. (Costantino et al. 2020) produced new Hi-C and ChIP-seq data, in order to better understand the relationship between the identified patterns of chromatin domains (information derived from Hi-C data analysis) and the cohesin residency regions (information derived from ChIP-seq data analysis). They used cells arrested in mitosis, and studied both the wild type strain and mutants altered for cohesin or its regulators. Notably, to finely describe chromatin loop formation, they used a recent improvement of the Hi-C technique, called Micro-C XL (Hsieh et al. 2016). This method has the advantage of greatly improving the detection of short-range interactions (at the scale of nucleosomes), while still allowing the detection of whole genome chromatin interactions. This is of particular interest for a species like *S. cerevisiae*, in which chromosomes are very small compared to the other species (*S. cerevisiae* chromosomes range from 0.23 to 1.53 Mb only, see **Figure 2A**). As a result, they showed that in *S. cerevisiae* mitotic cells, high residency cohesin anchor regions (named “CARs” and detected with ChIP-seq) correspond to the genome-wide boundaries of chromatin loops (named “CAR domains” and detected with Micro-C XL). This was an important observation which *i*) supports the idea that the yeast genome is organized into recurrent defined chromatin structures delimited by cohesin and *ii*) that this spatial organization of chromosomes can impact genome functions, as in mammals. Still, in their original article, the authors only present Hi-C contact heatmaps to support their interpretations.

To go further in this and enrich visualization, we collected their Micro-C XL data and reanalyzed them with 3DGB, to create an updated 3D model of the spatial organization of the *S. cerevisiae* chromosomes (see **Methods**). Considering the very high precision of the data, we expected to observe both the general configuration (chromosome territories and Rabl conformation) and the fine chromatin domains (coiling), by zooming in and out with visualization software. Our results are presented in **Figure 3**. Note that the overall model (**Figure 3A**) is the same as the one shown in **Figure 2B**, but with different (color-blind friendly) colors used to identify the 16 chromosomes of the *S. cerevisiae* genome. ChIP-seq data were also reanalyzed (see **Methods**) to locate the CARs on the new 3D model. They are shown in yellow in **Figure 3**. As expected from Hi-C contact maps presented in the original paper of Costantino et al. (Costantino et al. 2020), we observed a continuous distribution of CARs along the *S. cerevisiae* chromosomes (**Figure 3A**). To complete this observation, we zoomed in on the chromosomes individually. An example of chromosome 13 is shown **Figure 3B**. Notably, we could observe that CARs, which are still represented by yellow beads on the 3D models (**Figure 3D**), were aligned in space, and this, independently of the length of the chromatin loops whose boundaries they represent (**Figure 3C**). This observation was true for all chromosomes (another example can be found in **Supplementary Figure S3**). 3D modeling of Hi-C contacts is therefore of significant interest here, revealing the existence of a cohesin skeleton, ensuring structural stability of the spatial organization of the *S. cerevisiae* chromosomes.

**Figure 3:**
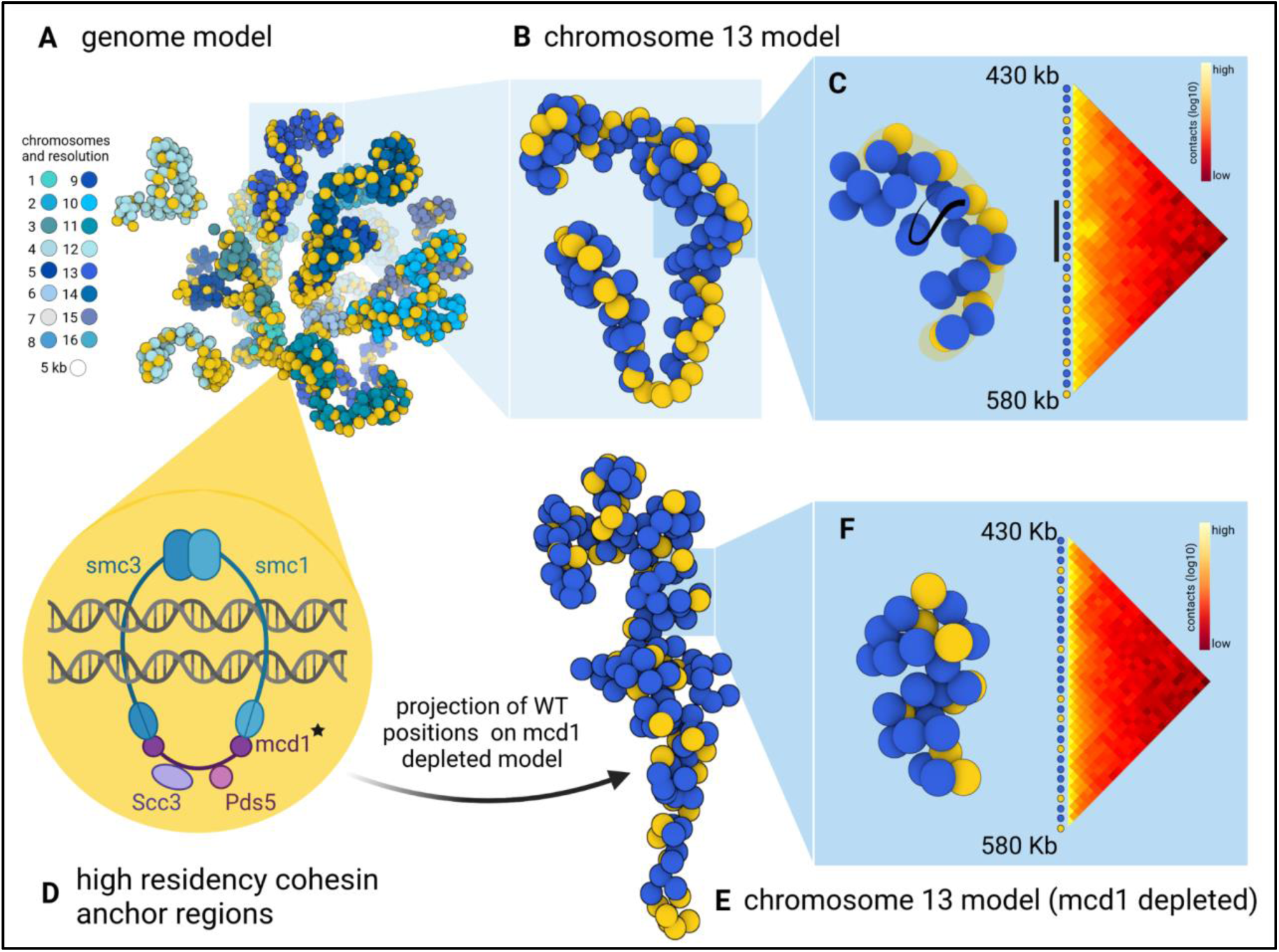
Revealing the 3D skeleton formed by cohesin in S. cerevisiae chromatin. **(A)** 3D model of S. cerevisiae chromatin, obtained with 3DGB, using the Micro-C XL data from Costantino et al. (Costantino et al. 2020). Each bead represents a 5 kb chromosomal region. Yellow color indicates genomic regions with a high value of ChIP-seq signal intensity of the Mcd1p cohesin subunit, in mitotically arrested cells. **(B)** Isolation of chromosome 13 from the overall structure, showing the spatial alignment of the DNA binding sites of cohesin. Cohesin structures a chromosomal skeleton on which chromatin loops can form. **(C)** Juxtaposition of the 3D model and the associated contact map, for the beads which are located between 430 and 580 kb. On the 3D structure, the cohesin backbone is highlighted in yellow. Viewing this spatial alignment of cohesin is complementary to the information contained in the contact map, i.e. the frequency of contacts is very high inside genomic regions delimited by yellow beads. **(D)** Schematic representation of the cohesin complex in S. cerevisiae. The protein Mcd1p, depleted in cells used to create the model shown in (E) and (F), is indicated with a black star. Genomic regions, which are attached by cohesin are referred to as CARs, i.e. high residency cohesin anchor regions, and are visualized in yellow on the 3D models. **(E)** Isolation of chromosome 13 from the overall structure, obtained with the mcd1 mutant (see the main text). The yellow beads correspond to the cohesive backbone as it was originally identified in the wild type (B). **(F)** Juxtaposition of the 3D model and the associated contact map, for the beads which are located between 430 and 580 kb. This is the same genomic region exposed in (C).

To verify the relevance of this observation, we also reanalyzed the Hi-C data obtained with an *S. cerevisiae* strain in which the gene encoding the subunit Mcd1 of the cohesin complex was depleted (**Figure 3D**, black star). Results are presented in **Figure 3E**. This time, the 3D models showed more irregular coiling of chromosomes and globule-like structures. Again, this observation is relevant to the original article of Costantino et al. in which the authors found that in the absence of cohesin, the mitotic chromatin is not entirely disorganized, but rather structured into globular domains, dependent on histone modifications (Costantino et al. 2020). An important question then was what happens, in this context, to the cohesin skeleton previously observed in the wild type strain. We therefore mapped the genomic positions of CARs, as they were defined based on the ChIP-seq data in the wild type strain, on the 3D “mutant” model (**Figure 3E**, yellow beads) and observed a complete destabilization of the cohesin backbone initially detected in wild type.

In summary, we illustrate here the interest of placing known 2D patterns of chromatin loops in a 3D context. Our observations reinforce the idea that a cohesin backbone stabilizes the spatial organization of chromosomes in *S. cerevisiae*. 3DGB brings a new perspective to the visualization compared to Hi-C contact maps: the flat loops are arranged in a coil supported by a backbone of aligned CAR regions.

##### Example #2: Oscillations in coiling of chromosome arms during the cell cycle in *S. pombe*

In the previous section, we were able to observe details of *S. cerevisiae* chromosome organization by zooming into the 3D model we built with 3DGB. This model was associated with cells arrested in mitosis and therefore represents a static view of the chromosomal organization at a very particular point in the yeast cell cycle. The goal of our second example was to evaluate the possibility of seeing, with 3D models, changes in the organization of chromosomes during the cell cycle. Indeed, chromosomes are subject to major structural constraints and undergo rearrangements at the different stages of the cell cycle (G2, M, G1 and S). We chose to explore this question using the data of Tanizawa et al., published in 2017 (Tanizawa et al. 2017). The authors applied an *in situ* Hi-C protocol to follow the organization of the fission yeast *S. pombe* genome throughout the cell cycle. The authors observed that during mitosis, chromosomes are structured into large (300 kb to 1 Mb) and small (30 - 40 kb) domains, which are respectively structured by condensin and cohesin protein complexes (Tanizawa et al. 2017). Based exclusively on their interpretation of Hi-C contact maps, they showed that if the mitotic organization into large domains gradually dissolves across the cell cycle, small domains remain relatively stable. They also hypothesized that the Rabl conformation was stable in interphase but disrupted during mitosis. With this dataset, we assessed our ability to observe these structural features in the 3D models obtained with 3DGB, at different stages of the *S. pombe* cell cycle. We collected Hi-C data from the original article (**Table 1**) and produced 3D models (see **Methods**), showing the organization of chromosomes at different time points of the S. pombe life cycle, after an initial synchronization of cells in phase G2. Our results are presented in **Figure 4**.

**Figure 4:**
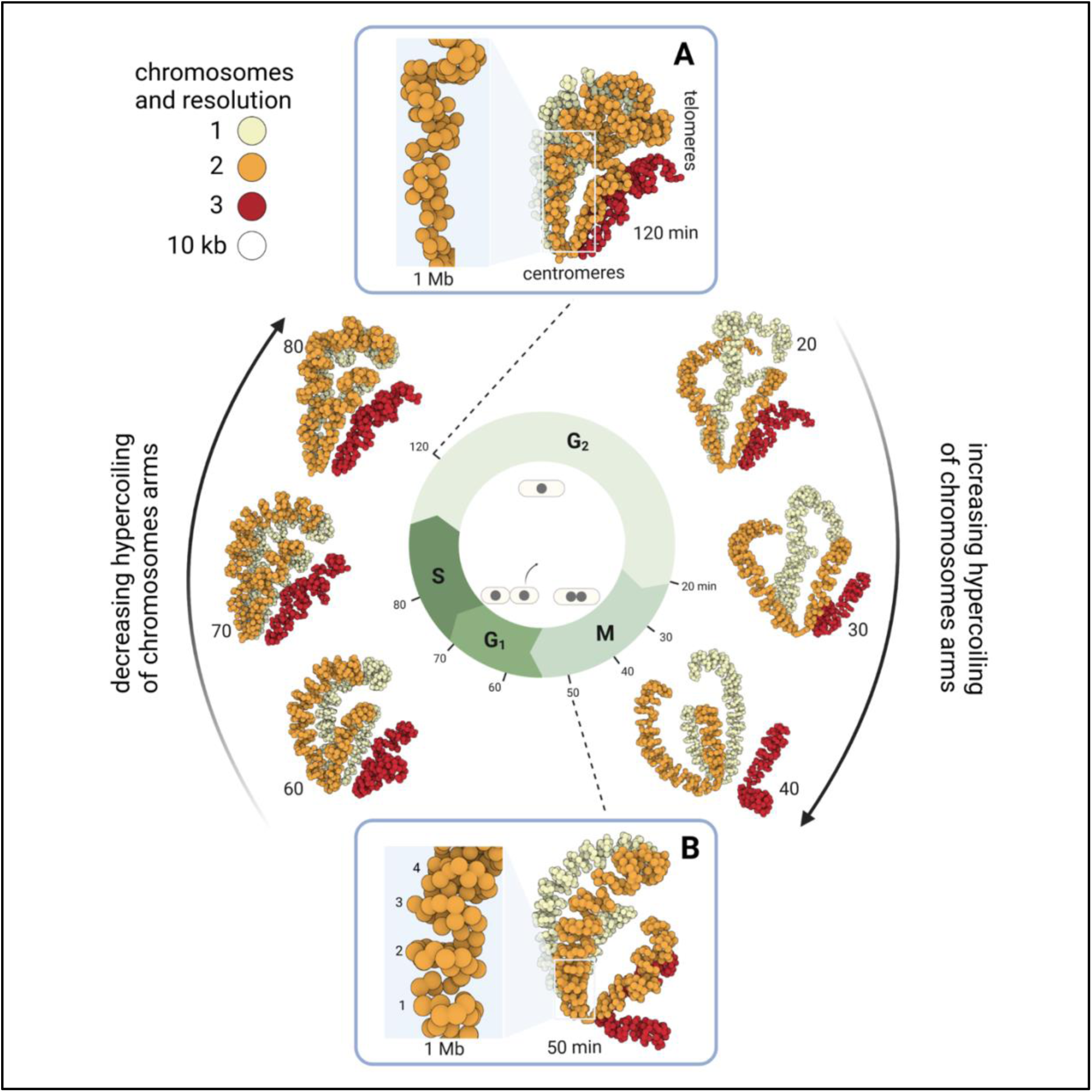
Dynamics of the spatial organization of S. pombe chromosomes during the cell cycle. 3D models obtained with 3DGB using Hi-C data from Tanizawa et al. (Tanizawa et al. 2017)*. Each bead represents a 10 kb chromosomal region and is colored according to the chromosome to which they belong (from 1 to 3). Eight models are shown here. They correspond to different organization of* S. pombe *chromosomes during the fission yeast cell cycle. After the synchronization of cells in phase G2, the following time points were analyzed: 20, 30, 40, 50, 60, 70, 80 and 120 minutes. Their correspondence with the cell cycle phases is shown in the middle. Two time points are thus in the G2 phase (20 and 120 minutes), three time points are in the M phase (30, 40 and 50 minutes), two time points are in the G1 phase (60 and 70 minutes), and finally one time point is in the S phase. Centromere and telomere regions are annotated. Centromeres of all chromosomes are constantly clustered, whatever the cell cycle time point. This is consistent with previous observations (see the main text). Coiling of chromosome arms, as described in the literature* (Mizuguchi et al. 2014; Tanizawa et al. 2017)*, is visible on the 3D models. Zooms are provided in (A) and (B) boxes, which correspond to time points (respectively 120 and 50 minutes) for which the intensity of coiling is minimal and maximal (see the main text for further explanation)*.

As expected, the Rabl conformation is visible, particularly during interphase (G1, S and G2 models, between 60 and 120 minutes). However, it becomes somewhat disorganized at the transition between G2 and Mitosis (**Figure 4**, 20 min model), even though the cluster of centromeres remains stable during the entire cell cycle. Telomeres of chromosomes 1 (yellow color) and 2 (orange color) also remain associated, whereas chromosome 3 (red color), in which ribosomal DNA repeats are found, occupied an external position consistent with compartmentalization of rDNA in the nucleolus. We also observed that the level of chromatin compaction varies over the cell cycle, with an observed minimum in G2 (**Figure 4A**) and an observed maximum in late mitosis (**Figure 4B**). Indeed, the 120 minute model shows uncoiled chromosome arms, with loose globule-like folding (**Figure 4A**) whereas the 50 minute model chromosome arms are structured into regular coils of ∼250 kb (**Figure 4B**). This more compact (see the zoom-in 1 Mb region **Figure 4A-B**) and structured (regular coiling) organization can be seen gradually appearing on the 20, 30 and 40 minute models (**Figure 4**, right arrow). Finally, it is important to keep in mind that the Hi-C data of each model were obtained independently, reinforcing the biological relevance of the observed similarities in relative position of chromosomes and level of compaction. Therefore, the highly organized mitotic structure gradually fades though the G1 and S phases to return to the Rabl conformation (**Figure 4**, left arrow).

In summary, the 3D models obtained with 3DGB consistently illustrate the oscillations in coiling of chromosome arms during the cell cycle in *S. pombe*. This strengthens the original study by Tanizawa et al. and opens new perspectives for further analysis, for example omics data integration.

##### Example #3: Massive relocalisation of histone marks in *N. crassa* heterochromatin regulator mutants

In the *N. crassa* genome, heterochromatic regions are a major component of the chromosome conformation (Galazka et al. 2016). Constitutive and facultative heterochromatin are respectively genomic regions that contain few genes with little transcription (constitutive heterochromatin) and genomic regions that contain genes with regulated gene repression (facultative heterochromatin). At the molecular level, constitutive and facultative heterochromatin can be distinguished by the presence of H3K9me3 or H3K27me2/3 histone marks (Galazka et al. 2016). Interactions between constitutive and facultative heterochromatin are complex and an important subject of discussion in the literature. In that respect, an emblematic observation is that the reduction of H3K9me3 in constitutive heterochromatin, causes the redistribution of H3K27me2/3. In particular, in a genetic context in which the heterochromatin protein 1 (Hp1, which recognize H3K9me3) is lost, H3K27me3 is depleted from facultative heterochromatin and H3K27me2 is gained at constitutive heterochromatin (Jamieson et al. 2016). Our objective was to assess the ability to see this phenomenon at the scale of the complete genome. To do so, we started by creating 3D models with 3DGB, from the wild type (WT) and HP1-deficient (*hpo*) strains. These models were then used for visual integration of ChIP-seq data, referring to the genomic location of histones with H3K9me3 and H3K27me2/3 post translational modifications (see **Methods**). Results are shown **Figure 5A**.

**Figure 5:**
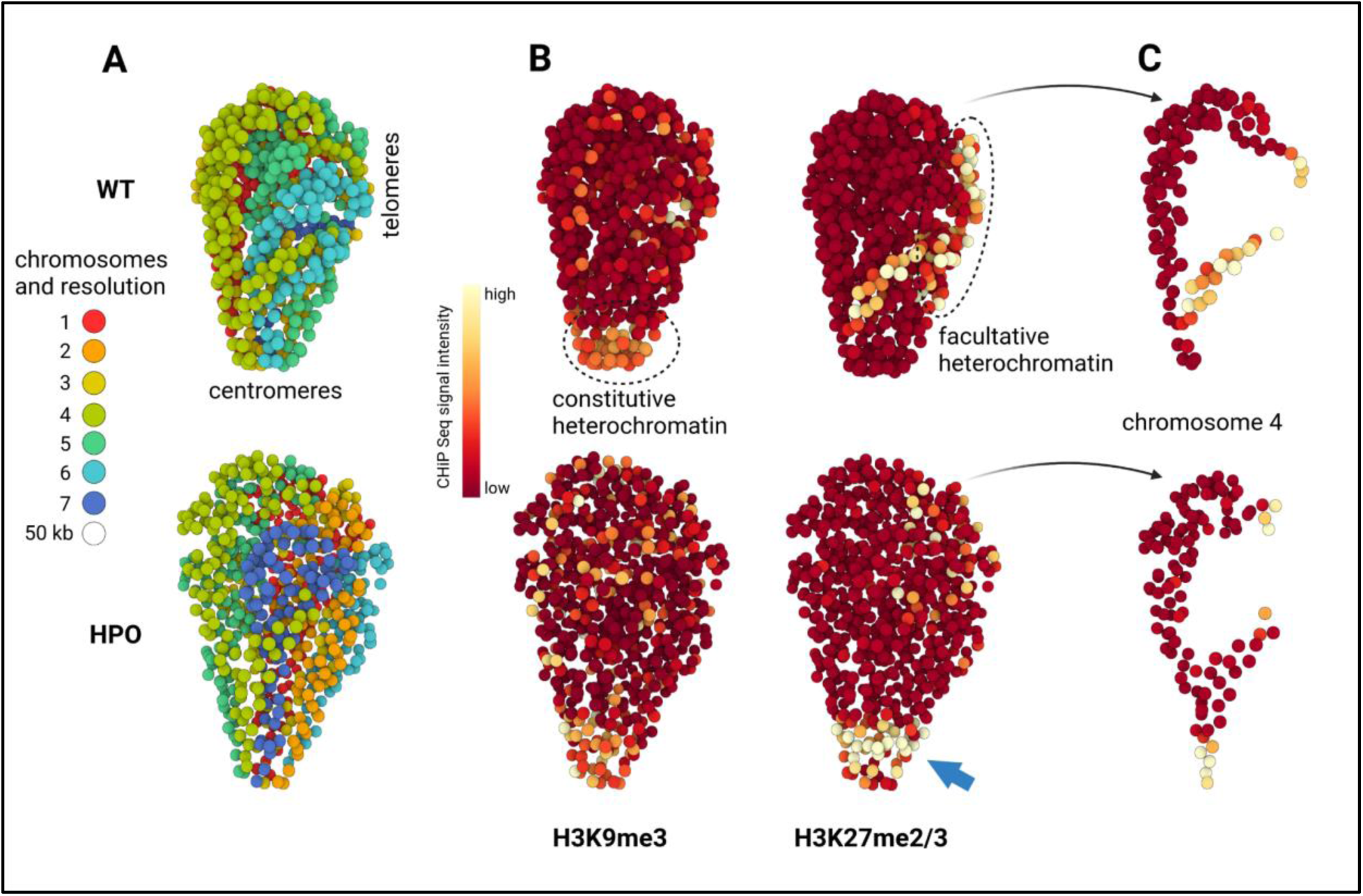
Visual integration of ChiP-seq data using 3D models of the N. crassa genome. (A) 3D models obtained with 3DGB using Hi-C data from Galaska et al. (Galazka et al. 2016)*, respectively in the wild type (WT) and HP1-deficient (hpo) strains (see* ***Table 1*** *and **Methods** section for technical details). Each bead represents a 50 kb chromosomal region and is colored according to the chromosome to which it belongs (from 1 to 7). The colocalization of centromeres and telomeres is observed in both models (centromeres on the bottom and telomeres on the right). **(B)** Same models as shown in (A), using the ChiP-seq signal intensity as a color code for beads (see **Methods**). ChIP-seq data are from Basenko et al.* (Basenko et al. 2015) *and refer to the genomic location of histones with H3K9me3 and H3K27me2/3 post translational modifications, respectively. In the WT model, the intensity of the ChIP-seq signal for H3K9me3 is particularly high (yellow color) around centromeres (compared to the rest of the genome, dark red color), whereas the intensity of the ChIP-seq signal for H3K27me3 is particularly high at telomeres. These observations are relevant to known functions of constitutive and facultative heterochromatin (see the main text). In the hpo strain, important changes in the intensities of ChIP-seq signals are observed with, notably, a massive relocalization of H3K27me3 high intensity signal to the centromeres (blue arrow). **(C)** Isolation of chromosome 4, better highlighting the changes in H3K27me2/3 ChIP-seq signal intensity between WT and hpo strains*.

As expected from the literature, we observed a slight difference between the two model organizations, essentially the intensity of chromatin compaction and the relative position of chromosomes 6 and 7 (**Figure 5A**). To extend this observation, we calculated for each 50 kb region represented by a bead in the 3D models, the intensity of ChIP-seq signals arising from experiments targeting H3K9me3 and H3K27me2/3 post-translational histone modifications (see **Methods**). Results are shown **Figure 5B** (whole genome) and **Figure 5C** (chromosome 4). In WT, we could observe as expected an accumulation of H3K9me3 histones in centromeres, relevant to constitutive heterochromatin, and an accumulation of H3K27me2/3 histones in sub-telomeric regions, relevant to facultative heterochromatin. In *hpo*, ChIP-seq signals appeared to be quite disorganized, especially for H3K27me2/3 marks, and massively relocalized to centromeres (blue arrow, **Figure 5B**). This observation was expected according to the literature, but for the first time it can be seen clearly, in a simple and integrated way.

This example illustrates the interest of 3D models in the context of multi-omics data integration. If Hi-C data allows us to better understand the organization of chromosomes, other omics data (such as ChIP-seq here) bring insights on the mechanisms of genome function. It is important to keep in mind that the images presented abstract a very large quantity of data, *i.e.* several million measurements (frequencies of contacts, probability of interactions, quality values, etc.). These 3D models can confirm or invalidate hypotheses on the overall functioning of genomes, and possibly lead to the formulation of new hypotheses.

#### Additional benefits of using 3DGB

##### Detection of inconsistencies between the spatial organization of chromosomes derived from Hi-C experiments and the genomic sequence used as reference

In the context of our analyses of Hi-C data in *N. crassa*, we faced an unexpected situation with respect to the reference genome sequence for this organism. Indeed, the *N. crassa* genome assembly, originally published in 2003 (Galagan et al. 2003), is composed of 7 chromosomes and 13 supercontigs, all available in GenBank under the accession “assembly nc12”. In 2016, Galazka et al. (Galazka et al. 2016) identified an inversion of the contig named “12.304”, corresponding to a large region in chromosome 6. In 2022, Rodriguez et al. (Rodriguez et al. 2022) further improved the original assembly “nc12” based on Hi-C data analysis. The updated assembly that integrates the improvements of Galazka et al. and Rodriguez et al. is available in the GEO database under the accession “assembly nc14”. When we started using 3DGB to create a 3D model of the wild-type *N. crassa* chromosome (**Figure 2**), we used as reference genome the “nc12” assembly, available from the GenBank database. Surprisingly, we observed in our inferred 3D model, an inconsistency in the order of the genomic regions, with respect to the succession of our beads (**Figure 6**). Our 3D model thus directly highlighted the inverted contig 12.304 on the chromosome 6, originally found by Galazka et al. At 10 kb resolution, the 95 beads used to represent this region were placed in the 3D space according to the information of Hi-C contacts only, independently of the reference genome assembly. This explains why we were able to highlight a mismatch between the chaining of beads in the 3D models and the numbering of these beads according to the genome sequence (**Figure 6A**). Note that when using the nc14 assembly of the genome as reference, the obtained 3D model of chromosome 6 we obtained was coherent both spatially (order of the beads) and sequentially (order of the genomics regions). From this experience, we developed an automated procedure (integrated to 3GDB as an option) to detect and correct inverted contigs by comparing the chaining of beads in the 3D model (based on the distance between adjacent beads, **Figure 6B**) and the numbering from the genomic sequence written in the FASTA file. This method also produces a corrected version of the 3D model and of the genome assembly (**Figure 6C**).

**Figure 6:**
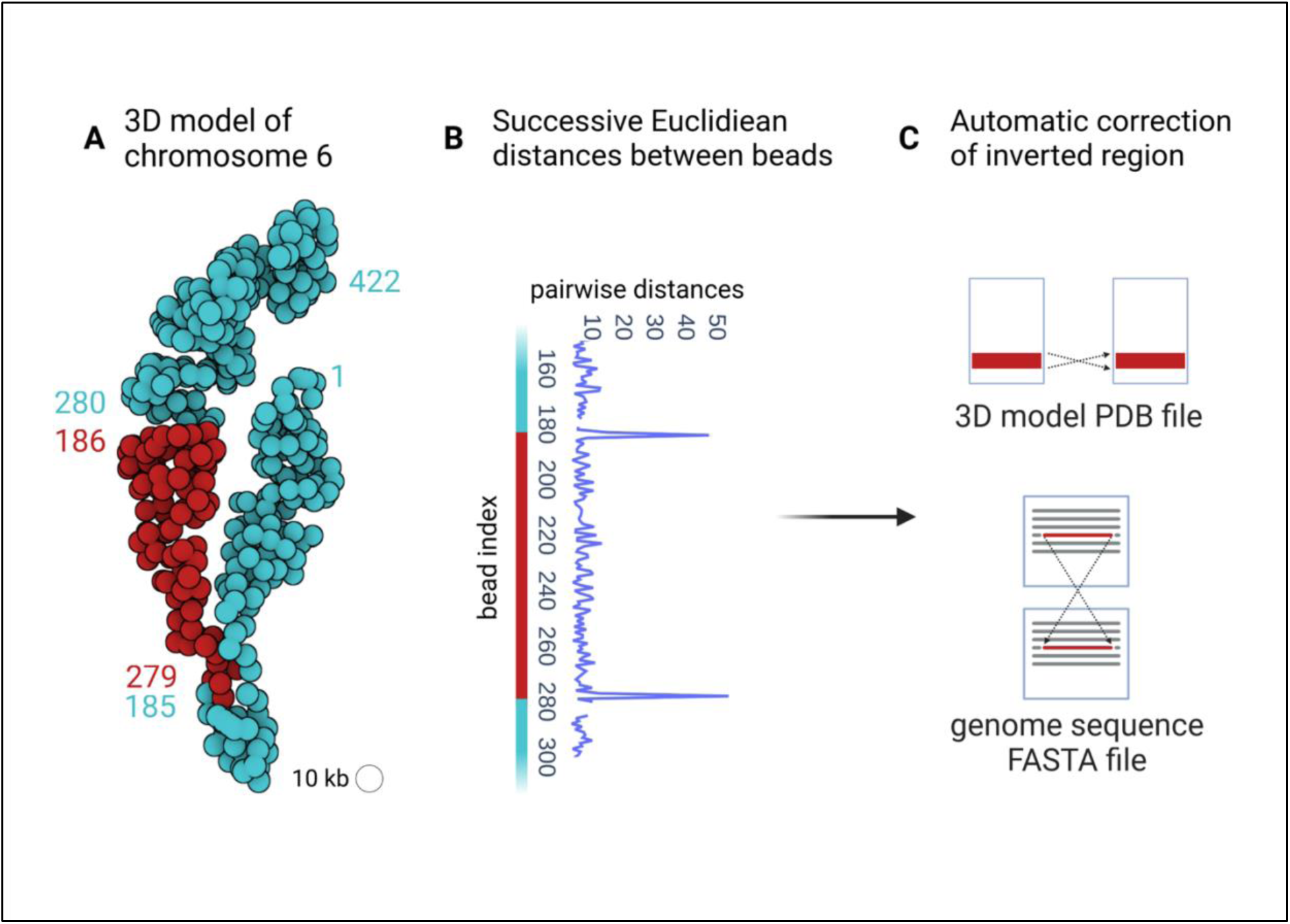
Inconsistency between the spatial organization of chromosome 6 in N. crassa and the reference genome assembly “nc12”. ***(A)** 3D model of chromosome 6 obtained with 3DGB using Hi-C data from Galaska et al. and the “nc12” assembly of the* N. crassa *genomic sequence as reference. Each bead represents a 10 kb chromosomal region. The numbers label the beads, according to the genomic region (DNA sequence) with which they are associated. Therefore, the bead number 1 corresponds to the interval]0, 10] kb of the linear genomic sequence, the bead number 2 corresponds to the interval]10, 20] kb, the bead number 3 corresponds to the interval]20, 30] kb, etc. Unexpected transitions between the pairs of beads 185 - 279 and 186 - 280 are shown with different colors (blue and red). **(B)** Euclidean distances between 3D beads with successive numbers as defined with genomic intervals on the DNA sequence. Unexpected high values are observed between bead pairs 185 - 186 and 279 - 280. They are in line with the structure shown in (A). **(C)** Based on the inconsistency revealed in the Euclidean distances, it is possible in 3DGB to automatically correct the genomic sequence used as a reference*.

##### Quantification of the stability of 3D models to random noise with multiple uses of 3DGB

The advantage of using a workflow like 3DGB is the possibility of automating tasks. In this part, we assess the stability of the 3D organization of the chromosomes with respect to potential inaccuracies in the contact frequencies measured with Hi-C (see **Methods**). Our strategy was to alter the original values of contact frequencies (issued from the Hi-C data) by adding a “shot noise” on the contact frequencies, and then run 3DGB to create an associated 3D model. If the intensity of the shot noise was low, the output model was expected to be close to the original model (obtained from the original data). The resulting RMSD score, calculated by comparing the 3D structure inferred from the reference *N. crassa* contact map and the 3D structure inferred from the altered *N. crassa* map, was also expected to be low. Altogether, we assessed 23 levels of intensity of the shot noise (see **Methods**) and generated, automatically, a total of 1150 models. Our results are summarized in **Supplementary Figure S5**. As expected, we observed an increase in the RMSD scores, when the intensity of the shot noise increases. This was more striking for the global structure (**Supplementary Figure 5B**, green boxplots) than for individual chromosomes (**Supplementary Figure 5B**, blue boxplots). This observation underlines the greater sensitivity of the organization of the chromosomal territories compared to the internal organization of the individual chromosomes.

## Discussion

The objective of this work was to explore the interest of creating 3D visualizations of chromosome organization to complement the classic contact maps producing during Hi-C data analysis. We found that, although methods to create 3D models exist, they are not often exploited, and this is particularly true of fungal genome studies. The emblematic 3D representation of the complete genome of *S. cerevisiae* had been obtained more than 10 years ago, but only partial structures were generated for *S. pombe* and *N. crassa* (see **Supplementary Figure S1**). Our first intention was therefore to create, from existing bioinformatics tools and Hi-C datasets, complete models for these three species. Unlike what has been done for *S. cerevisiae* by Duan et al. (Duan et al. 2010), we wanted to build models without specifying initial physical constraints regarding chromosome positioning. The Pastis-NB method was particularly suitable for this task. To automate the generation of 3D models, we built 3DGB, a high-throughput Hi-C data processing workflow (**Figure 1**). With this workflow, we provided enriched PDB files for advanced visualization with molecular viewer software (**Figure 2**) and we created several hundred models in reasonable computing times (**Supplementary Figure S5**). The 3DGB workflow is available to other scientists who would like to add 3D models to their analyses of Hi-C data.

The relevance of 3D models of the spatial organization of chromosomes is an important question of our study. Beyond the production of a “beautiful image”, what is the added value of these models for Hi-C data analysts? It is indeed important to keep in mind that these models are not “pictures” of the interior of cell nuclei. They are based on experimental data, whose quality may vary and significantly impact the inferred model. Ultimately, they represent measurements of contact frequencies observed at the scale of cell populations and therefore, they are simplified representations of reality. This work on fungal genomes taught us that the visualization of 3D models is profitable for several reasons. First, it gives an overall representation of the organization of the chromosomes (**Figure 2**). While contact maps are very useful for identifying local chromatin structures (at the scale of kilo base pairs), 3D models allow a complete zoom out, at the scale of the entire chromosome, giving a global representation of all contact frequencies. Second, 3D models can both serve as the final visualization of the results of a Hi-C study, but also as a starting point for further data integration. The results we presented for the *S. cerevisiae* and *S. pombe* genomes are examples of final visualization results. On the one hand, our models revealed important features of the organization of chromosomes, respectively a backbone of cohesin (**Figure 3**) and hypercoiling dynamics (**Figure 4**), which are difficult to fully understand from the contact heatmaps alone. On the other hand, the results we presented for *N. crassa* are examples of original data integration (**Figure 5**). This time, 3D models are starting points for additional exploration. The coloring of the 3D representation of the models, based on quantitative measurements from “omics” experiments, albeit a simple idea, is particularly effective for studying global phenomena, such as the massive relocalization of histone marks in a mutant, as shown here for *N. crassa*.

Still, it is worth considering that contact-based modeling of the genome 3D organization has some limits. They remain a 3D representation of a Hi-C based contact network, with chromatin regions modeled as independent beads, each representing several kilo base pairs. The sequencing depth of the Hi-C data and the chosen resolution for their analysis have a direct impact on the accuracy of the final model: for a given number of reads, the lower the resolution, the fewer the inter chromosomal contacts are detected. We observed that when the sequencing depth is low, the chromosomes in the 3D models are more distant from each other, for example in the mitotic *S. cerevisiae* and *S. pombe* models (**Figure 2**). In such a situation, it is difficult to discriminate between the biological compaction of the chromatin and the limitations due to the Hi-C experimental technique, when only looking at the final volume occupied by a model. However, structural features like level of coiling are still visible (**Figures 3** and **4**) and can provide insight at the level of the biological compaction of chromatin, compensating for technical limitations (resulting in chromosomes which are more spaced). As stated previously, another inherent limitation of 3D models is that they are “static”, “population averaged” pictures of a biological system (chromatin organization) known to be highly dynamic and variable from cell to cell. These artifacts will exist as long as the Hi-C data are snapshots obtained from cell populations. Still, it is important to note that Rabl conformations and chromosome territories were observed in all species (**Figure 2**), even with the *N. crassa* model, which is an average representation derived from Hi-C experiments performed on an asynchronous cell population. Therefore, with synchronous cells, the situation can be expected to be even better. This is what we observed with *S. pombe* models. Because the cells were initially synchronized, the Hi-C datasets produced at different stages of the cell cycle revealed several other interesting genome organizations in the inferred 3D models (**Figure 4**) and we thus managed to render the dynamics of chromatin. As for the “population average” limit, single cell Hi-C strategies are developing (Stevens et al. 2017), opening the promising perspective of creating 3D genome models of single cells. A final limit of current 3D models is the missing regions of the fungal genomes. As illustrations, the models presented here are built using genomic sequences in which the rDNA repeats were deleted, thus preventing the correct representation of structural features in the nucleolus (explaining the hole we can observe on chromosome 12 in *S. cerevisiae* models, **Figures 2** and **3**). An additional important simplification arises from the joint representation of sister chromatids and the complexity of working with diploid genomes (in classical Hi-C data analyses, the contact measurements cannot be distinguished between identical sequences). Strategies are emerging to solve haplotypes (Oomen et al. 2020; Mitter et al. 2020) and the software used in 3DGB has tuned options for this: at the mapping step, HiC-Pro can build allele-specific contact maps if SNP information is provided. At the 3D modeling stage, the Poisson-based method of Pastis has been extended to haplotype resolution. Even if this allows us to imagine creation of more complete models in the future, it is important to keep in mind that the fungal genomes examined here, *i)* remain haploid during the cell cycle and *ii)* have sister chromatids tightly maintained together by cohesion. We therefore believe that representation of fungal chromosomes as one string of beads is informative.

Overall, the limits underline the dependence of 3D modeling on the quality of experimental Hi-C raw data and generation methods. We are confident however, that the technical improvements that are being introduced at a rapid pace will progressively minimize these limitations. Importantly, even though the technical ceiling is not yet reached, we have already managed to highlight important fungal genome characteristics in a novel way by reanalyzing public datasets. For *N. crassa*, the 6 3D models produced by 3DGB condensed the information of 5 research articles and about 20 raw FASTQ data files into one rich large-scale illustration (**Figure 5**). They bring new opportunities for visual integration of omics data: while Hi-C analysis often implies a “zoom-in” logic, focusing on precise regions of a contact map, 3D modeling completes and enhances this logic with a “zoomed-out” vision.

In conclusion, we have presented a holistic approach that is favorable to intuition and new hypotheses. We explored large-scale integration of epigenetic ChIP-seq data into the 3D context of chromatin, but any suitable combination of omics datasets can be made using the genome model as a visual support. For example, we are currently exploring the interpolation of genes on the 3D genome structure. 3DGB could be adapted to provide a model at the gene (one bead each) level, to allow the mapping of gene-dependent omics data, such as RNA-seq, on the chromosome structure. Gene positions need not be accurate, but rather provide a convenient way to plot omics data on top of the 3D organization of the chromatin. Going even further, it could be of interest to integrate information from protein occupancy or transcription factor interaction networks. Seeing chromatin domains as loops of a coil instead of distinct entities could open new insights on inter-domain gene regulation.

## Methods

### 3DGB implementation and workflow management

#### Technical details

3DGB was orchestrated with the open-source workflow management system [Snakemake](https://snakemake.readthedocs.io/en/stable/) (Mölder et al. 2021), which automates the different steps of a data analysis in a human-readable, Python-based language. 3DGB is structured in multiple individual rules depicted in **Supplementary Figure S1**. Software environments were deployed and isolated in a Conda environment (for Pastis and all custom Python scripts) and a Singularity container (for HiC-Pro).

#### 3DGB inputs

3DGB requires the following inputs: Hi-C FASTQ files (provided as SRA IDs or as local FASTQ files), a reference genome sequence provided in a FASTA file (without mitochondrial DNA), one or several enzyme restriction site motifs, and finally, values for the output resolution to draw the contact map and generate 3D models.

#### Main steps in the analysis

The first steps of the workflow format the necessary information for the read mapping step: *(i)* FASTQ files are downloaded and compressed if not already provided by the user; *(ii)* a list of fragments derived from the genome FASTA file and the enzyme restriction site motif is generated by HiC-Pro (Servant et al. 2015) version 3.1.0; *(iii)* the size of each chromosome from the genome FASTA file is extracted with a custom script and *(iv)* the reference genome is indexed with Bowtie2 version 2.4.4. Next, HiC-Pro generates a set of ‘valid pairs’, which are relevant reads used to generate and normalize contact frequencies (or counts). For a given resolution, these values are then used to compute contact maps (heatmaps) and to serve as inputs for Pastis (Varoquaux et al. 2014, 2021) (version as of July, 21st, 2021) to build a consensus 3D model. In 3DGB, the Pastis-NB method is used. It iteratively computes 3D models of the organization of chromosomes, through negative binomial contact count modelization. For better reproducibility and portability, HiC-Pro is used in a Singularity image provided by the authors of the software. Pastis is installed in a conda environment. In 3DGB, a final step to refine the 3D model provided by Pastis is available. It consists in predicting the coordinates for additional beads, for which initial calculations were missing in the 3D model from Pastis. Missing coordinates usually occur in centromeric and telomeric regions. Predictions are performed by interpolation, using monotonic cubic splines, as implemented by the pchip method in the Scipy Python library (Virtanen et al. 2020). Note that beads with missing coordinates localized at the extremities of chromosomes are discarded and beads with aberrant coordinates are filtered out based on a threshold applied to the Euclidean distance value calculated between neighboring beads.

#### 3DGB outputs

3D models are stored in PDB files. From the raw model produced by Pastis, 3DGB annotates chromosomes by numbers, based on the reference sequence specified in the 3DGB inputs. Examples of models, fully annotated with 3DGB, are provided as PDB files in Supplementary Data (see source code and data availability). In PDB files, the chromosome annotation is present in the residue id (1, 2, 3…), in the chain id (A, B, C…) and in the residue name (C01, C02, C03…). This adaptation of the standard residue and chain fields in PDB files is particularly useful, allowing immediate visualization of chromosomes with visualization software. Eventually, quantitative values can be associated with the beads of the 3D model (for instance ChIP-Seq signal intensity), using the B factor field in PDB files. Most PDB viewers can display B factors on top of the structures. This is the case for Mol*, used to create images presented in this article.

### Analyses of experimental datasets in fungal species

#### Access to raw data from SRA database

All the Hi-C seq, Micro-C XL seq and ChIP seq raw FASTQ files were downloaded from the SRA database, see **Table 1** and **Supplementary Table 1** (second tab for ChIP seq) for all SRA ids. The reference genomes of *S. cerevisiae* and *S. pombe* were obtained from the NCBI Genome database, respectively version R64 (S288C) and ASM294v2.19. The nc14 version of the *N. crassa* genome was downloaded from the supplementary data of Rodriguez et al. on NCBI GEO database, GEO dataset GSE173593. FASTA files with chromosome sequences only were generated as 3DGB input.

#### Application of 3DGB to create 3D models

3DGB was applied with default parameters (FASTQ files, see **Table 1** for all details) and configuration files can be accessed in Zenodo and Github. For *S. cerevisiae*, 2 models were generated at 5 kb resolution from 4 FASTQ files (total of 31 028 357 valid pairs) and ChiP-Seq information from 2 FASTQ files were integrated. For *S. pombe*, 8 models at 10 kb resolution were built, from 19 FASTQ files (total of 136 321 555 valid pairs). For *N. crassa*, 3 models were created at 50 kb and 10 kb resolutions, from 14 FASTQ files (total of 115 588 951 valid pairs) and ChiP-Seq information from 9 FASTQ files were integrated. The chosen resolutions of the models were defined according to the technique used (Hi-C or Micro-C XL), the number of reads and their quality.

#### Evaluation of 3D model stability to random noise

The *hpo* mutant *N. crassa* contact frequency matrix was used as a “reference” to generate “noisy contact frequency matrices”. The *hpo* mutant was selected because it has the lowest number of valid read pairs (2,922,526) and this sample is the most likely to be sensitive to noise. Let *x_ij_* be the number of counts in the contact frequency matrix of this sample, with *i* and *j* the indices of the beads ranging from 1 to *n* = 825, with *n* the total number of beads for this sample. To create an “altered” or “noisy” *N. crassa* contact frequency matrix, it was set *y_ij_* = *x_ij_* + *x_ij_* the number of counts in the noisy contact frequency matrix, where *e_ij_* is a value sampled from a Poisson distribution with parameter λ, independently for each pair of beads (*i*, *j*). Note that the contract frequency matrices are symmetric and *e_ij_* = *e_ji_* for each pair of beads. The Poisson distribution was used, because it models shot noise, *i.e.* the variability in read counts that is due to the read sampling process rather than to biological variations (Anders and Huber 2010). The level of noise (*i.e.* the mean parameter λ of the Poisson distribution used to generate shot noise) varies among the following values 0.1, 5, 10, 15, 20 up to 110. For each value of λ, 50 replicates were generated, *i.e.* 50 noisy contact frequency matrices. To assess the stability of 3D structures, the RMSD were computed between the 3D structure inferred from the reference *N. crassa* contact frequency matrix (the raw contact matrix with counts *x_ij_*) and the 3D structure inferred from the altered *N. crassa* maps (the noisy contact frequency matrix with counts *y_ij_*). This procedure leads to 50 values of RMSD for each value of λ. The **Supplementary Figure S5** represents the RMSD values for λ ranging from 0.1 to 50. Above the value λ = 50, the RMSD values reach a threshold and do not increase with the value of λ. The RMSD between each one of the seven chromosomes of *N. crassa* were also computed, for each value of λ and each of the 50 replicates.

#### Visual integration of omics data

The same pipeline was used for integration of ChIP-Seq data from *N. crassa* and *S. cerevisiae*. To perform alignment, Bowtie2 was used with default parameters (**Supplementary Table 1, tab 2)**. The program samtools was used to sort and index the SAM files into bam files, and the program bamCoverage was used to generate 5 kb or 50 kb .bedgraph files with RPKM normalization (for *S. cerevisiae* and *N. crassa* respectively). Each bin in the bedgraph corresponds to one bead of the 3D model. Note that the resolution of the bedgraph and of the 3D model are the same, allowing the integration of the data into the PDB file. For *S. cerevisiae*, the threshold was set to 80 counts after normalization, converting the continuous ChIP-seq signal into a binary signal highlighting the high residency CARs (Costantino et al. 2020). For *N. crassa*, the ChIP-seq data was kept continuous to highlight the epigenetic mark distribution. The values below 20 counts are discarded for visualization.

## Data access

The 3DGB workflow is available on GitHub: https://github.com/data-fun/3d-genome-builder. The .PDB files of the 13 models presented in this study are available on Zenodo <u>10.5281/zenodo.7740302</u>, as well as animated GIF for 4 of them. The .YML (config files) for generating those 13 models are available on GitHub and on Zenodo.

## Competing interest statement

The authors declare that they have no competing interests.

## Acknowledgments

We thank Nelle Varoquaux for useful discussions. This work was funded by the Agence Nationale pour la Recherche (MINOMICS project, Grant Number ANR-19-CE45-0017). Figures were made on BioRender.

---

**Supplementary Figure S1:**
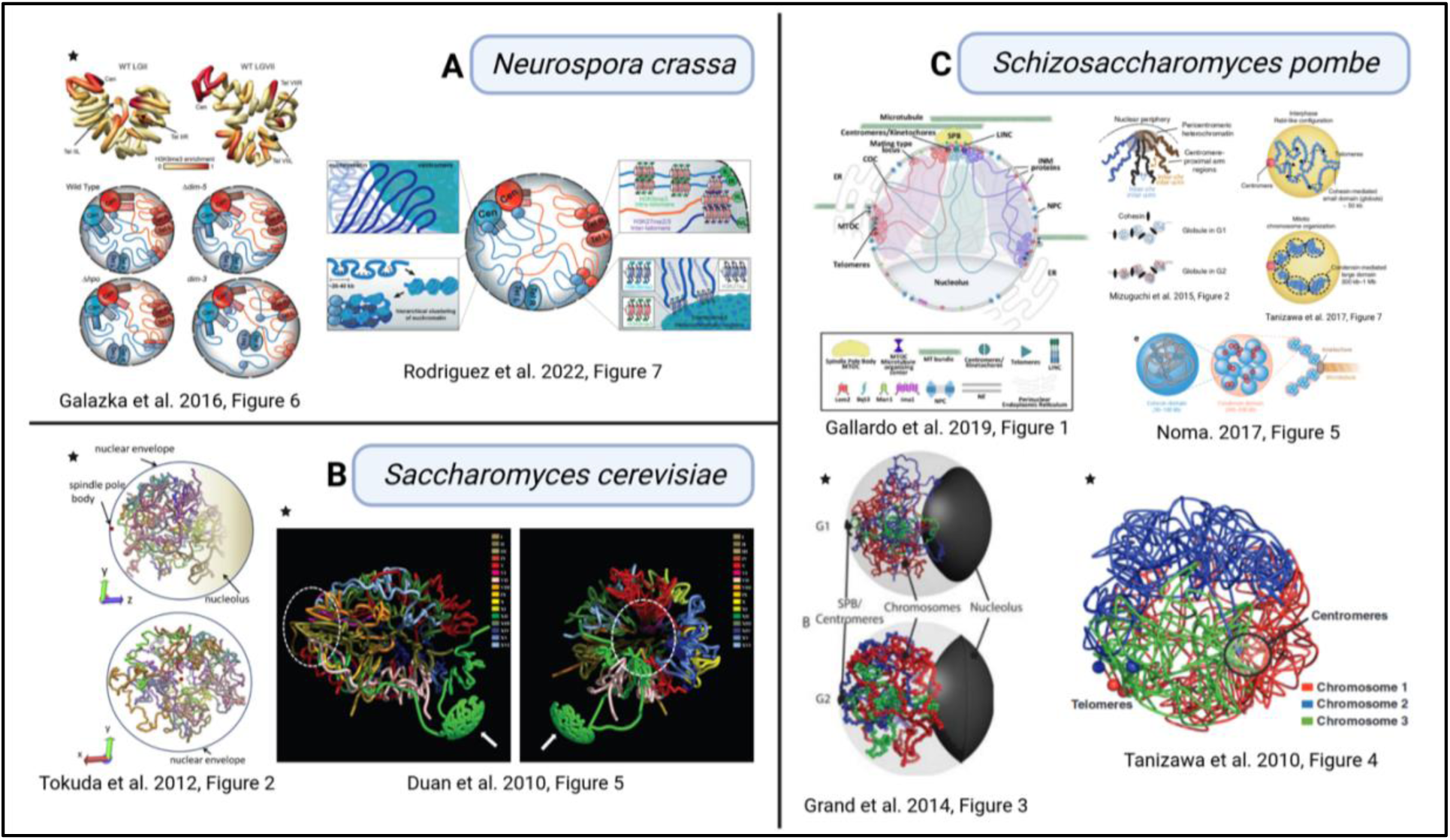
Spatial organization of fungal chromosomes, as described in the literature. 3D models of Neurospora crassa (A), Saccharomyces cerevisiae (B) and Schizosaccharomyces pombe (C) genomes, from several previously published articles. Notably, these models are either drawings, meaning that they are interpretations of the data made by the authors, or real 3D objects (see black stars), meaning that they arise from the application of dedicated algorithms for the calculation of spatial coordinates, based on Hi-C contact measurements. In both situations, these models summarize current knowledge of the overall organization of genomes of these three species of fungi (see the main text for descriptions of interesting properties).

**Supplementary Figure S2:**
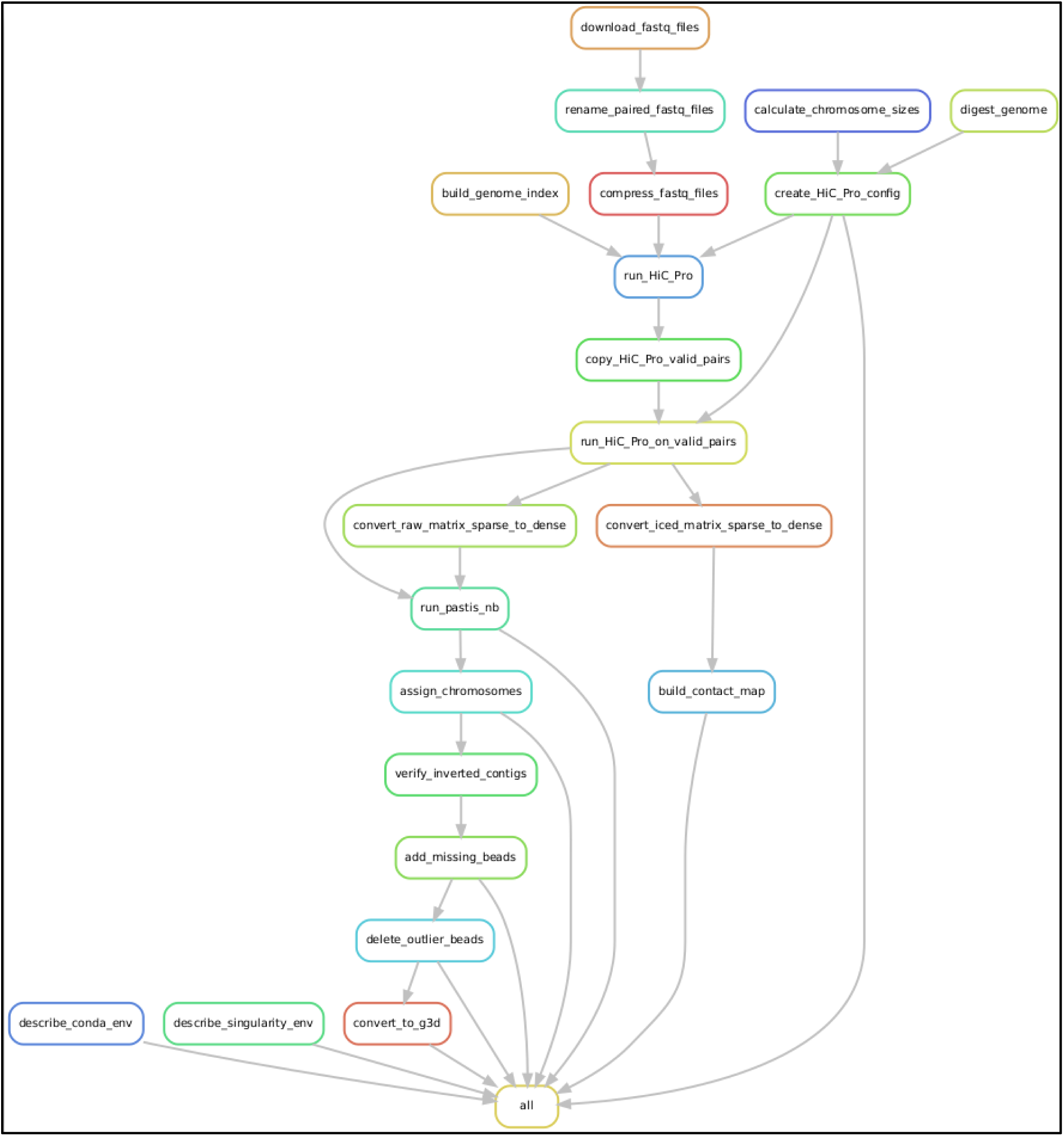
**Directed acyclic graph of the 3DGB workflow as exported from Snakemake**. Each box refers to a Snakemake rule with defined inputs and outputs. Rules can be run in parallel by Snakemake. More information can be found in https://github.com/data-fun/3d-genome-builder.

**Supplementary Figure S3:**
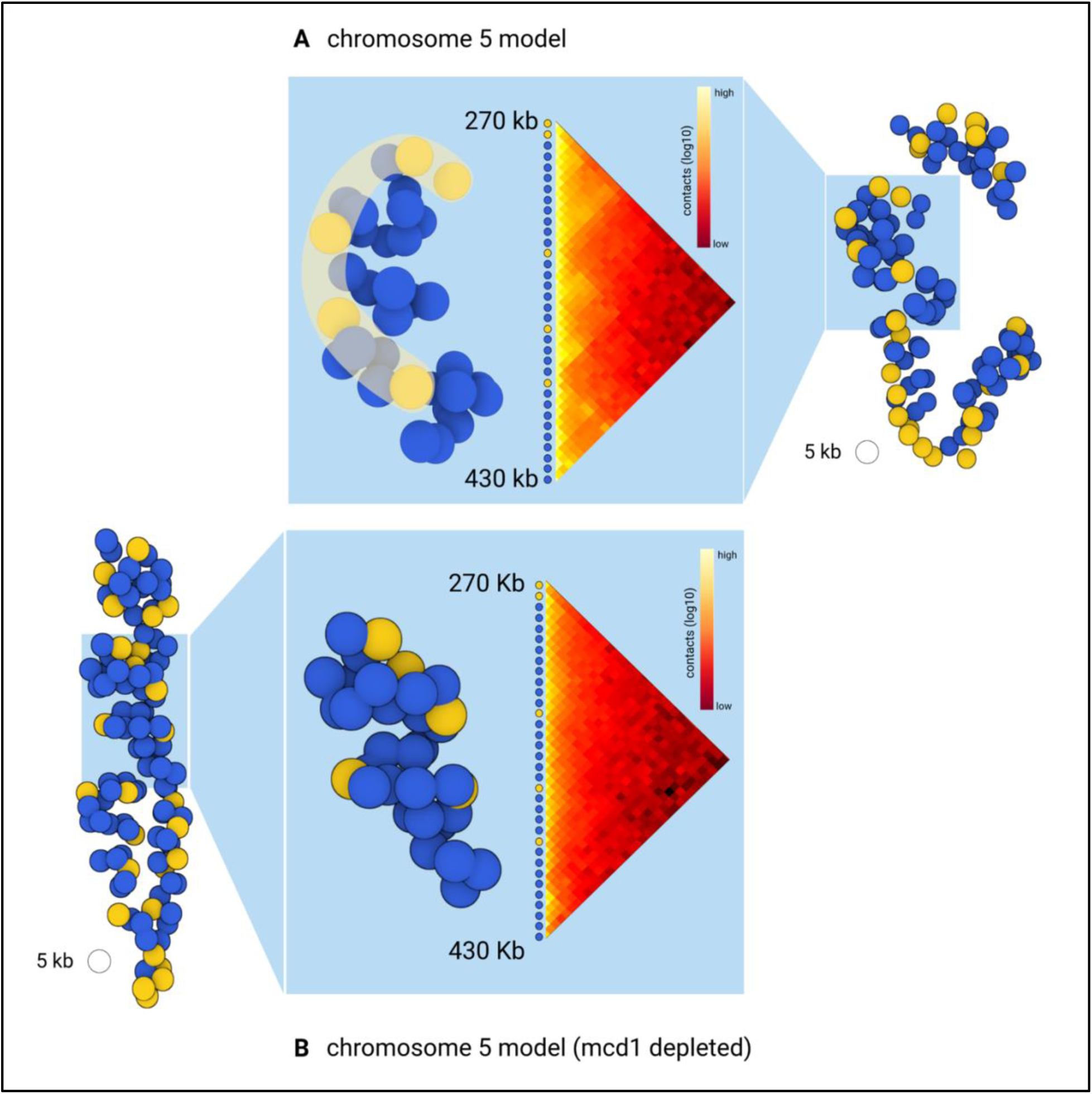
3D skeleton formed by cohesin in S. cerevisiae chromatin, example of chromosome 5. (A) Isolation of chromosome 5 from the overall structure obtained with the wild type strain and **(B)** isolation of chromosome 5 from the overall structure obtained with the strain depleted for the Mcd1 cohesin subunit. Yellow beads correspond to the backbone of CARs (see the main text), as defined in the wild type situation.

**Supplementary Figure S5:**
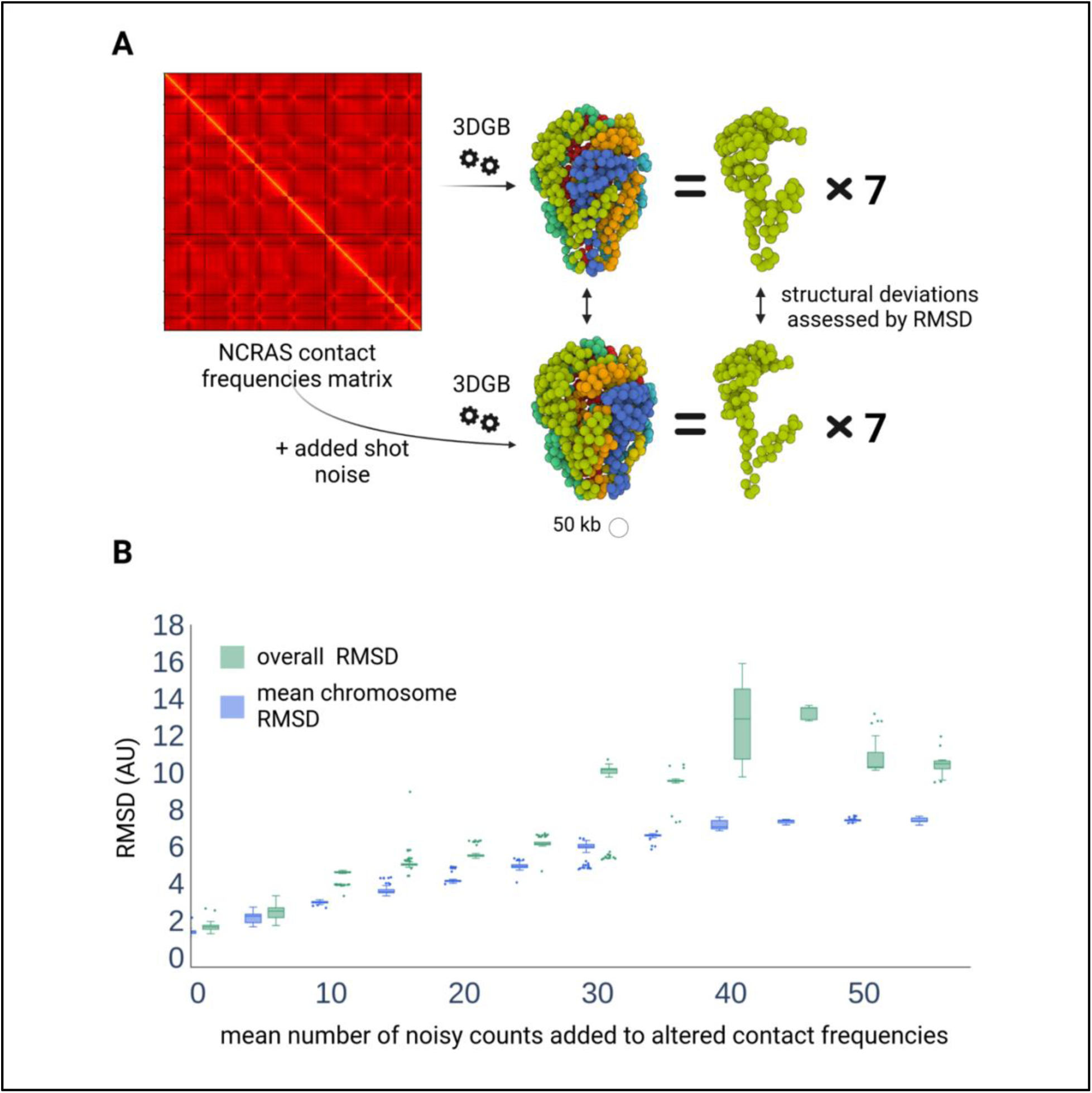
Quantification of the stability of 3D models to random noise added to original contact frequencies. (A) Each pairwise frequency of contacts is randomly modified by adding a count value drawn from a Poisson distribution with a specified mean parameter (see Methods). 3D models are created with 3DGB (at 50 kb resolution) and compared by calculating RMSD scores. (B) Values of RMSD scores (arbitrary unit) obtained for different values of noise, i.e. different values for the mean parameter used in the Poisson distribution. For a given noise value, 50 models are generated, the RMSD score is either calculated on the overall structure composed of all chromosomes (green boxes) or averaged on the RMSD scores obtained for the 7 individual chromosomes (blue boxes).

## Notes

https://doi.org/10.5281/zenodo.7740302

https://github.com/data-fun/3d-genome-builder

### Competing Interest Statement

The authors have declared no competing interest.

## References

1. Anders S, Huber W. 2010. Differential expression analysis for sequence count data.

2. Arifulin EA, Musinova YR, Vassetzky YS, Sheval EV. 2018. Mobility of Nuclear Components and Genome Functioning. Biochemistry Moscow 83: 690–700.

3. Asbury TM, Mitman M, Tang J, Zheng WJ. 2010. Genome3D: A viewer-model framework for integrating and visualizing multi-scale epigenomic information within a three-dimensional genome. BMC Bioinformatics 11: 444.

4. Basenko EY, Sasaki T, Ji L, Prybol CJ, Burckhardt RM, Schmitz RJ, Lewis ZA. 2015. Genome-wide redistribution of H3K27me3 is linked to genotoxic stress and defective growth. Proc Natl Acad Sci USA 112: E6339–E6348.

5. Bauer CR, Hartl TA, Bosco G. 2012. Condensin II Promotes the Formation of Chromosome Territories by Inducing Axial Compaction of Polyploid Interphase Chromosomes ed. R.S. Hawley. PLoS Genet 8: e1002873.

6. Baumeister W. 2022. Cryo-electron tomography: The power of seeing the whole picture. Biochemical and Biophysical Research Communications 633: 26–28.

7. Carlier F, Li M, Maroc L, Debuchy R, Souaid C, Noordermeer D, Grognet P, Malagnac F. 2021. Loss of EZH2-like or SU(VAR)3–9-like proteins causes simultaneous perturbations in H3K27 and H3K9 tri-methylation and associated developmental defects in the fungus Podospora anserina. Epigenetics & Chromatin 14: 22.

8. Costantino L, Hsieh T-HS, Lamothe R, Darzacq X, Koshland D. 2020. Cohesin residency determines chromatin loop patterns. eLife 9: e59889.

9. Cremer T, Cremer C. 2001. Chromosome territories, nuclear architecture and gene regulation in mammalian cells. Nat Rev Genet 2: 292–301.

10. Denecker T, Zhou Li Y, Fairhead C, Budin K, Camadro J-M, Bolotin-Fukuhara M, Angoulvant A, Lelandais G. 2020. Functional networks of co-expressed genes to explore iron homeostasis processes in the pathogenic yeast Candida glabrata. NAR Genomics and Bioinformatics 2: lqaa027.

11. Djekidel MN, Wang M, Zhang MQ, Gao J. 2017. HiC-3DViewer: a new tool to visualize Hi-C data in 3D space. Quant Biol 5: 183–190.

12. Duan Z, Andronescu M, Schutz K, McIlwain S, Kim YJ, Lee C, Shendure J, Fields S, Blau CA, Noble WS. 2010. A three-dimensional model of the yeast genome. Nature 465: 363–367.

13. Dundr M, Misteli T. 2001. Functional architecture in the cell nucleus.

14. Durand NC, Shamim MS, Machol I, Rao SSP, Huntley MH, Lander ES, Aiden EL. 2016. Juicer Provides a One-Click System for Analyzing Loop-Resolution Hi-C Experiments. Cell Systems 3: 95–98.

15. Fritz AJ, Sehgal N, Pliss A, Xu J, Berezney R. 2019. Chromosome territories and the global regulation of the genome. Genes Chromosomes Cancer 58: 407–426.

16. Galagan JE, Calvo SE, Borkovich KA, Selker EU, Read ND, Jaffe D, FitzHugh W, Ma L-J, Smirnov S, Purcell S, et al. 2003. The genome sequence of the filamentous fungus Neurospora crassa. Nature 422: 859–868.

17. Galazka JM, Klocko AD, Uesaka M, Honda S, Selker EU, Freitag M. 2016. Neurospora chromosomes are organized by blocks of importin alpha-dependent heterochromatin that are largely independent of H3K9me3. Genome Res 26: 1069–1080.

18. Gallardo P, Barrales RR, Daga RR, Salas-Pino S. 2019. Nuclear Mechanics in the Fission Yeast. Cells 8: 1285.

19. Goodsell DS. 2009. The machinery of life. 2nd ed., corrected. Copernicus Books, New York.

20. Grand RS, Pichugina T, Gehlen LR, Jones MB, Tsai P, Allison JR, Martienssen R, O’Sullivan JM. 2014. Chromosome conformation maps in fission yeast reveal cell cycle dependent sub nuclear structure. Nucleic Acids Research 42: 12585–12599.

21. Grognet P, Timpano H, Carlier F, Aït-Benkhali J, Berteaux-Lecellier V, Debuchy R, Bidard F, Malagnac F. 2019. A RID-like putative cytosine methyltransferase homologue controls sexual development in the fungus Podospora anserina ed. E. Gladyshev. PLoS Genet 15: e1008086.

22. Guido Van Rossum, Fred L. Drake. 2009. Python 3 Reference Manual. CreateSpace.

23. Hoencamp C, Dudchenko O, Elbatsh AMO, Brahmachari S, Raaijmakers JA. 2022. 3D genomics across the tree of life identifies condensin II as a determinant of architecture type.

24. Hsieh T-HS, Fudenberg G, Goloborodko A, Rando OJ. 2016. Micro-C XL: assaying chromosome conformation from the nucleosome to the entire genome. Nat Methods 13: 1009–1011.

25. Im W, Liang J, Olson A, Zhou H-X, Vajda S, Vakser IA. 2016. Challenges in structural approaches to cell modeling. Journal of Molecular Biology 428: 2943–2964.

26. Jamieson K, Wiles ET, McNaught KJ, Sidoli S, Leggett N, Shao Y, Garcia BA, Selker EU. 2016. Loss of HP1 causes depletion of H3K27me3 from facultative heterochromatin and gain of H3K27me2 at constitutive heterochromatin. Genome Res 26: 97–107.

27. Jensen EC. 2013. Overview of Live-Cell Imaging: Requirements and Methods Used. Anat Rec 296: 1–8.

28. Jerković I, Cavalli G. 2021. Understanding 3D genome organization by multidisciplinary methods. Nat Rev Mol Cell Biol. http://www.nature.com/articles/s41580-021-00362-w (Accessed June 18, 2021).

29. Kempfer R, Pombo A. 2020. Methods for mapping 3D chromosome architecture. Nat Rev Genet 21: 207–226.

30. Kim K-D, Tanizawa H, Iwasaki O, Noma K. 2016. Transcription factors mediate condensin recruitment and global chromosomal organization in fission yeast. Nat Genet 48: 1242– 1252.

31. Lelandais G, Remy D, Malagnac F, Grognet P. 2022. New insights into genome annotation in Podospora anserina through re-exploiting multiple RNA-seq data. BMC Genomics 23: 859.

32. Li D, Harrison JK, Purushotham D, Wang T. 2022. Exploring genomic data coupled with 3D chromatin structures using the WashU Epigenome Browser. Nat Methods 19: 909–910.

33. Li J, Zhang W, Li X. 2018. 3D Genome Reconstruction with ShRec3D+ and Hi-C Data. IEEE/ACM Trans Comput Biol and Bioinf 15: 460–468.

34. Love MI, Huber W, Anders S. 2014. Moderated estimation of fold change and dispersion for RNA-seq data with DESeq2. Genome Biol 15: 550.

35. Misteli T. 2020. The Self-Organizing Genome: Principles of Genome Architecture and Function. Cell 183: 28–45.

36. Mitter M, Gasser C, Takacs Z, Langer CCH, Tang W, Jessberger G, Beales CT, Neuner E, Ameres SL, Peters J-M, et al. 2020. Conformation of sister chromatids in the replicated human genome. Nature 586: 139–144.

37. Mizuguchi T, Fudenberg G, Mehta S, Belton J-M, Taneja N, Folco HD, FitzGerald P, Dekker J, Mirny L, Barrowman J, et al. 2014. Cohesin-dependent globules and heterochromatin shape 3D genome architecture in S. pombe. Nature 516: 432–435.

38. Mölder F, Jablonski KP, Letcher B, Hall MB, Tomkins-Tinch CH, Sochat V, Forster J, Lee S, Twardziok SO, Kanitz A, et al. 2021. Sustainable data analysis with Snakemake. F1000Res 10: 33.

39. Nogales E, Scheres SHW. 2015. Cryo-EM: A Unique Tool for the Visualization of Macromolecular Complexity. Molecular Cell 58: 677–689.

40. Noma K. 2017. The Yeast Genomes in Three Dimensions: Mechanisms and Functions. Annu Rev Genet 51: 23–44.

41. Nowotny J, Wells A, Oluwadare O, Xu L, Cao R, Trieu T, He C, Cheng J. 2016. GMOL: An Interactive Tool for 3D Genome Structure Visualization. Sci Rep 6: 20802.

42. O’Donoghue SI. 2021. Grand Challenges in Bioinformatics Data Visualization. Front Bioinform 1: 669186.

43. Oluwadare O, Highsmith M, Cheng J. 2019. An Overview of Methods for Reconstructing 3-D Chromosome and Genome Structures from Hi-C Data. Biol Proced Online 21: 7.

44. Oomen ME, Hedger AK, Watts JK, Dekker J. 2020. Detecting chromatin interactions between and along sister chromatids with SisterC. Nat Methods 17: 1002–1009.

45. Poinsignon T, Gallopin M, Camadro J-M, Poulain P, Lelandais G. 2022. Additional insights into the organization of transcriptional regulatory modules based on a 3D model of the Saccharomyces cerevisiae genome. BMC Res Notes 15: 67.

46. Pouokam M, Cruz B, Burgess S, Segal MR, Vazquez M, Arsuaga J. 2019. The Rabl configuration limits topological entanglement of chromosomes in budding yeast. Sci Rep 9: 6795.

47. Radulović S, Sunkara S, Rachel R, Leitinger G. 2022. Three-dimensional SEM, TEM, and STEM for analysis of large-scale biological systems. Histochem Cell Biol 158: 203–211.

48. Razin SV, Borunova VV, Iarovaia OV, Vassetzky YS. 2014. Nuclear matrix and structural and functional compartmentalization of the eucaryotic cell nucleus. Biochemistry Moscow 79: 608–618.

49. Reckel S, Löhr F, Dötsch V. 2005. In-Cell NMR Spectroscopy. ChemBioChem 6: 1601–1606.

50. Rieber L, Mahony S. 2017. miniMDS: 3D structural inference from high-resolution Hi-C data. Bioinformatics 33: i261–i266.

51. Rodriguez S, Ward A, Reckard AT, Shtanko Y, Hull-Crew C, Klocko AD. 2022. The genome organization of *Neurospora crassa* at high resolution uncovers principles of fungal chromosome topology ed. J. Dekker. G3 Genes|Genomes|Genetics 12: jkac053.

52. Rodriguez-Granados NY, Ramirez-Prado JS, Veluchamy A, Latrasse D, Raynaud C, Crespi M, Ariel F, Benhamed M. 2016. Put your 3D glasses on: plant chromatin is on show. EXBOTJ 67: 3205–3221.

53. Sear RP, Pagonabarraga I, Flaus A. 2015. Life at the mesoscale: the self-organised cytoplasm and nucleoplasm. BMC Biophys 8: 4.

54. Sehnal D, Bittrich S, Deshpande M, Svobodová R, Berka K, Bazgier V, Velankar S, Burley SK, Koča J, Rose AS. 2021. Mol* Viewer: modern web app for 3D visualization and analysis of large biomolecular structures. Nucleic Acids Research 49: W431–W437.

55. Sénécaut N, Poulain P, Lignières L, Terrier S, Legros V, Chevreux G, Lelandais G, Camadro J-M. 2022. Quantitative Proteomics in Yeast: From bSLIM and Proteome Discoverer Outputs to Graphical Assessment of the Significance of Protein Quantification Scores. In *Yeast Functional Genomics* (ed. F. Devaux), Vol. 2477 of Methods in Molecular Biology, pp. 275–292, Springer US, New York, NY https://link.springer.com/10.1007/978-1-0716-2257-5_16 (Accessed March 3, 2023).

56. Servant N, Varoquaux N, Lajoie BR, Viara E, Chen C-J, Vert J-P, Heard E, Dekker J, Barillot E. 2015. HiC-Pro: an optimized and flexible pipeline for Hi-C data processing. Genome Biol 16: 259.

57. Smyth MS. 2000. x Ray crystallography. Molecular Pathology 53: 8–14.

58. Stevens TJ, Lando D, Basu S, Atkinson LP, Cao Y, Lee SF, Leeb M, Wohlfahrt KJ, Boucher W, O’Shaughnessy-Kirwan A, et al. 2017. 3D structures of individual mammalian genomes studied by single-cell Hi-C. Nature 544: 59–64.

59. Tanizawa H, Iwasaki O, Tanaka A, Capizzi JR, Wickramasinghe P, Lee M, Fu Z, Noma K. 2010. Mapping of long-range associations throughout the fission yeast genome reveals global genome organization linked to transcriptional regulation. Nucleic Acids Research 38: 8164–8177.

60. Tanizawa H, Kim K-D, Iwasaki O, Noma K. 2017. Architectural alterations of the fission yeast genome during the cell cycle. 31.

61. Todd S, Todd P, McGowan SJ, Hughes JR, Kakui Y, Leymarie FF, Latham W, Taylor S. 2021. CSynth: an interactive modelling and visualization tool for 3D chromatin structure ed. A. Valencia. Bioinformatics 37: 951–955.

62. Tokuda N, Terada TP, Sasai M. 2012. Dynamical Modeling of Three-Dimensional Genome Organization in Interphase Budding Yeast. Biophysical Journal 102: 296–304.

63. Trieu T, Oluwadare O, Wopata J, Cheng J. 2019. GenomeFlow: a comprehensive graphical tool for modeling and analyzing 3D genome structure ed. B. Berger. Bioinformatics 35: 1416–1418.

64. Varoquaux N, Ay F, Noble WS, Vert J-P. 2014. A statistical approach for inferring the 3D structure of the genome. Bioinformatics 30: i26–i33.

65. Varoquaux N, Noble WS, Vert J-P. 2021. Inference of genome 3D architecture by modeling overdispersion of Hi-C data. Bioinformatics http://biorxiv.org/lookup/doi/10.1101/2021.02.04.429864 (Accessed January 14, 2022).

66. Virtanen P, Gommers R, Oliphant TE, Haberland M, Reddy T, Cournapeau D, Burovski E, Peterson P, Weckesser W, Bright J, et al. 2020. SciPy 1.0: fundamental algorithms for scientific computing in Python. Nat Methods 17: 261–272.

67. Wong AMH, Eleftheriades GV. 2013. An Optical Super-Microscope for Far-field, Real-time Imaging Beyond the Diffraction Limit. Sci Rep 3: 1715.

68. Yardımcı GG, Noble WS. 2017. Software tools for visualizing Hi-C data. Genome Biol 18: 26.

69. Zhang C, Xu Z, Yang S, Sun G, Jia L, Zheng Z, Gu Q, Tao W, Cheng T, Li C, et al. 2020. tagHi-C Reveals 3D Chromatin Architecture Dynamics during Mouse Hematopoiesis. Cell Reports 32: 108206.

